# Benchmarking long-read RNA-sequencing technologies with *LongBench:* a cross-platform reference dataset profiling cancer cell lines with bulk and single-cell approaches

**DOI:** 10.1101/2025.09.11.675724

**Authors:** Yupei You, Ashleigh N. Solano, James Lancaster, Margaux David, Changqing Wang, Jia Wei Tan, Shian Su, Camilla Pasquali, Kathleen Zeglinski, Reza Ghamsari, Manveer Chauhan, Josie Gleeson, Yair D. J. Prawer, Jin Ng, Bénédicte Dubois, Isabelle Cleynen, Marie-Liesse Asselin-Labat, Nadia M. Davidson, Kate D. Sutherland, Michael B. Clark, Quentin Gouil, Matthew E. Ritchie

## Abstract

Long-read RNA sequencing enables full-length transcript profiling and improved isoform resolution, but variable platforms and evolving chemistries demand careful benchmarking for reliable application. We present *LongBench*, a matched, multi-platform reference dataset spanning bulk, single-cell, and single-nucleus transcriptomics across eight human lung cancer cell lines with synthetic spike-in controls. *LongBench* incorporates three state-of-the-art long-read protocols alongside Illumina short reads: Oxford Nanopore Technologies (ONT) PCR-cDNA, ONT direct RNA, and PacBio Kinnex. We systematically evaluate transcript capture, quantification accuracy, differential expression, isoform usage, variant detection, and allele-specific analyses. Our results show high concordance in gene-level differential analyses across protocols, but reduced consistency for transcript-level and isoform analyses due to length- and platform-dependent biases. Single-cell long-read data are highly concordant with bulk for high-confidence features, though single-nuclei data show reduced feature detection. *LongBench* provides one of the largest publicly available long-read benchmarking resources, enabling rigorous cross-platform evaluation and guiding technology selection for transcriptomic research.

---

Long-read RNA sequencing (RNA-seq) provides access to the still largely untapped complexity of transcriptomes, facilitating isoform discovery and quantification. These capabilities have successfully revealed transcript diversity and highlighted biologically and clinically relevant isoforms in human diseases, including cancer^(1–5)^ and neurological disorders^(6–9)^. Pacific Biosciences (PacBio)^(10)^ and Oxford Nanopore Technologies (ONT)^(11)^ are leading long-read platforms with distinct strategies. PacBio’s recently released Kinnex protocol^(12)^, a commercialised version of MAS-seq^(13)^, boosts throughput by concatenating cDNAs into extended circular inserts. ONT has introduced Kit 14 and RNA004 chemistry for cDNA and direct RNA sequencing respectively, which, along with improved basecalling models, have substantially increased read accuracy. Advances in instrumentation (e.g., PacBio Revio, ONT PromethION) have pushed cDNA yields to over 100 million reads per flow cell^(14)^, enabling high-throughput transcriptomics. ONT direct RNA sequencing also allows simultaneous detection of RNA base modifications, while avoiding reverse transcription and PCR biases associated with cDNA-based sequencing approaches^(15–18)^. Both technological and bioinformatics landscapes continue to evolve rapidly, highlighting the need for systematic benchmarking.

The rapid development and growth of both long-read RNA-seq and single-cell transcriptomics catalysed interest in developing long-read single-cell approaches, which have revealed the isoform information and genetic variants missed by standard short-read single-cell RNA-seq^(1,19,20)^. Early attempts to apply ONT and PacBio to single-cell RNA-seq faced key limitations in read accuracy and sequencing yield. These necessitated the creation of custom protocols to improve accuracy, such as R2C2 ^(21)^, or the use of matched short-read scRNA-seq to provide key information such as cell barcodes, cell expression profiles or cell-type identification^(1,19,22–27)^. With recent improvements in software, throughput and sequencing accuracy, ONT–only protocols without short-reads have become feasible^(20,28)^. Concurrently, the standardisation of PacBio “HiFi” sequencing and developments in PacBio single-cell RNA-seq protocols, including targeted enrichment of barcoded cDNA^(29)^ and improved read yield in Kinnex, have enhanced isoform recovery and sensitivity^(19,30,31)^. As single-cell long-read protocols evolve, an expanding ecosystem of computational tools has emerged to support tasks such as cell barcode demultiplexing, isoform annotation, and transcript quantification^(1,20,28,32–36)^. Nevertheless, knowledge of the comparative performance of different methods remains limited, highlighting the importance of comprehensive benchmarking to properly evaluate different methods.

Several foundational benchmarking studies have provided key insights into isoform detection, transcript quantification and platform-specific biases, yet are often constrained by study designs that limit their relevance to current workflows. Weirather et al. ^(37)^ found superior consensus read accuracy, lower error rates, and better splice site determination for PacBio, and higher throughput and comparable isoform detection for ONT. Soneson et al. ^(38)^ revealed protocol-specific transcript coverage biases with ONT cDNA and direct RNA. Dong et al. ^(39)^ compared ONT PromethION and Illumina, highlighting differences in isoform quantification and differential expression analyses. Larger studies, including LR-GASP^(40)^ and SG-NEx^(41)^, demonstrated variability in isoform discovery, improved major isoform quantification, and enhanced transcript diversity with long reads. More recently, Wissel et al. ^(42)^ reported more reliable transcript-level differential expression of PacBio Kinnex over Illumina, and Zajac et al. ^(43)^ and Scoones et al. ^(44)^ highlighted advances in single-cell long-read analyses. Together, these studies have established essential groundwork for long-read RNA-seq benchmarking. However, notable limitations persist: (1) most benchmarks evaluate only a subset of platforms (Supplementary Fig. 1A); (2) many rely on synthetic or homogeneous inputs with minimal biological replication (Supplementary Fig. 1B), restricting generalisability; and (3) due to the fast-moving nature of this field, most feature obsolete or outdated sequencing chemistries or instrumentation. To support future transcriptomic research, comprehensive bench-marking resources that incorporate current platforms, biological diversity, and multiple data modalities are needed.

Here we present *LongBench*, a long-read transcriptomic dataset for benchmarking sample preparation workflows, sequencing technologies and analysis methods. *LongBench* includes 3 modalities (bulk, single-cell and single-nucleus transcriptomics) across 3 long-read sequencing technologies (ONT PCR-cDNA and PacBio Kinnex for all modalities and ONT direct RNA for bulk). The experimental design included samples from 8 lung cancer cell lines representing 3 distinct disease groups and spiked-in transcripts. A diverse panel of cancer cell lines was included to capture realistic biological variation with all samples sequenced deeply. These design features facilitated systematic assessment of transcript capture efficiency and quantification accuracy across transcripts of varying lengths and abundances, as well as differential expression testing between disease groups. The single-cell and single-nucleus components enable transcript-level analyses at cell resolution, while also allowing assessment of the impact of additional library preparation and computational steps. By providing a comprehensive resource for com-paring sequencing platforms and analysis tools across diverse conditions, this dataset aims to guide researchers in making informed decisions and advancing the use of long-read RNA-seq in transcriptomic research.

## Results

### Experimental design with realistic levels of biological variation

*LongBench* includes eight lung cancer cell lines derived from patients representing three distinct cancer types (Fig.1A): Lung Adenocarcinoma (LUAD) and two Small Cell Lung Cancer (SCLC) subtypes, SCLC-A and SCLC-P, distinguished by transcriptional regulators, ASCL1 and POU2F3 ^(45)^. Synthetic RNA spike-ins, namely Sequins^(46)^ (mix A or B) and Lexogen SIRV set 4, were added to each bulk RNA sample to provide ground truth internal standards (Supplementary Table 1). A total of 32 bulk RNA-seq libraries were generated using four sequencing protocols: ONT PCR-cDNA, ONT direct RNA (dRNA), PacBio Kinnex cDNA, and Illumina short-read sequencing. In parallel, a pooled mixture of cells from all eight cell lines was prepared for single-cell and single-nucleus RNA-seq. cDNA libraries were generated with 10X Genomics Chromium and sequenced with ONT PCR-cDNA, PacBio Kinnex cDNA, and Illumina, yielding three scRNA-seq and three snRNA-seq datasets. Together, these bulk, single-cell, and single-nucleus datasets enable a comprehensive cross-platform comparison. Our short-read sequencing data validated the design and cell line selection by demonstrating clustering by cancer type while preserving realistic within-group biological variability (Fig.1B-C), supporting the reliability and biological relevance of subsequent differential analysis. With *LongBench*, we present an overview of multiple downstream analyses (Fig.1D), detailed in subsequent sections.

**Fig. 1.**
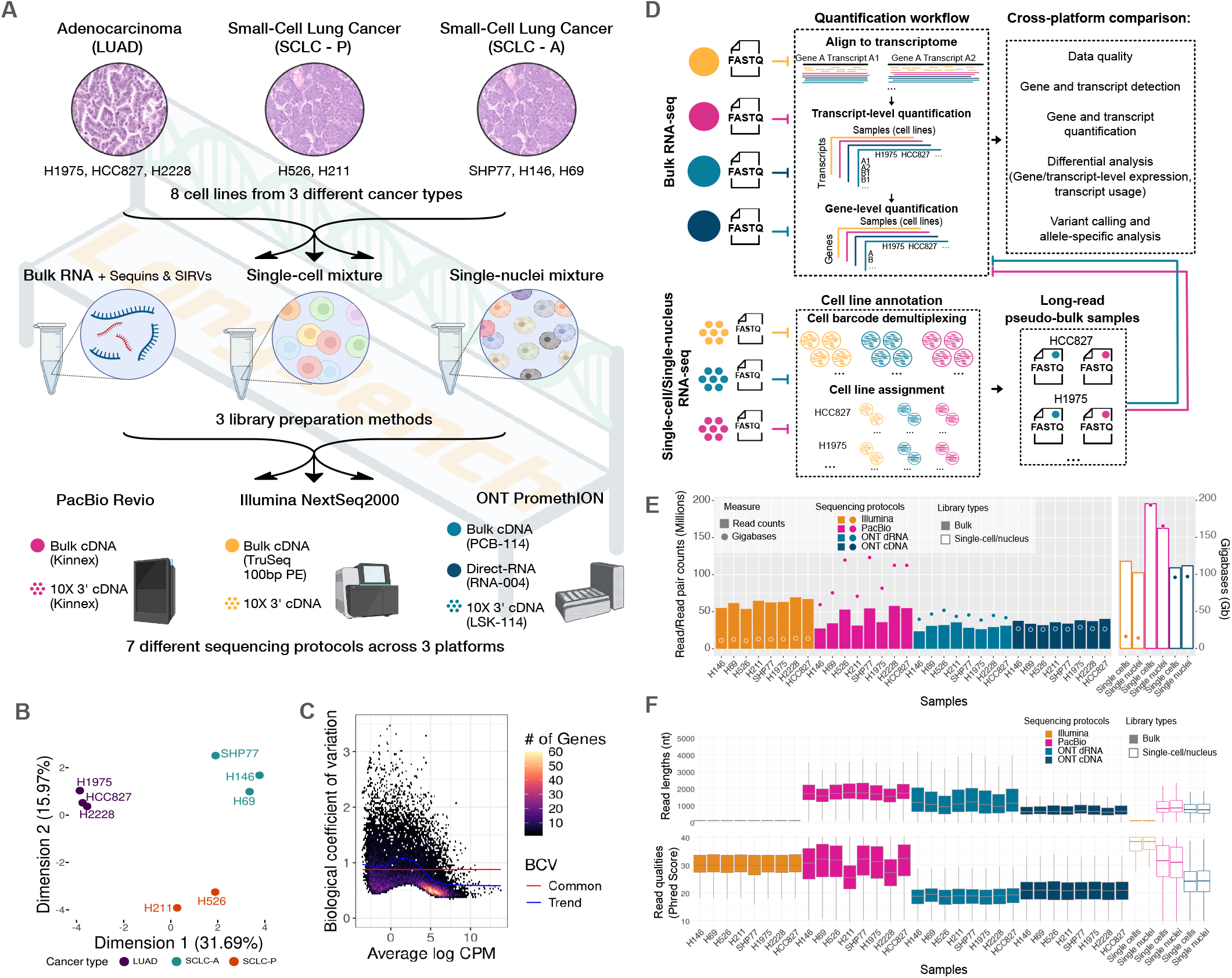
*LongBench* study design and data overview. **A**. Design of the *LongBench* study, which included bulk, single-cell and single-nucleus samples derived from 8 lung cancer cell lines, profiled across 3 long-read sequencing technologies, including Oxford Nanopore Technologies PCR-cDNA and PacBio Kinnex for all modalities and Oxford Nanopore Technologies direct RNA sequencing for bulk, as well as Illumina short-read cDNA sequencing across all sample types. **B**. Multidimensional scaling (MDS) plot of samples based on the gene quantification matrix from short-read RNA-seq dataset in *LongBench*. **C**. Biological coefficient of variation (BCV) generated from the short-read RNA-seq dataset using data from the 8 cancer cell lines. The BCV was estimated using edgeR. **D**. Summary of the data available and the main analyses performed in this study. Colours indicate the different sequencing technologies, consistent with those in A. **E**. The number of reads and bases generated for each sample. **F**. A summary of read lengths and qualities from each library. Note that the read lengths for Illumina were fixed to 101bp (bulk) and 90bp (single-cell/single-nuclei).

### *LongBench* features high-quality, deeply sequenced RNA-seq data

*LongBench* comprises 2.17 billion total reads, including 1.45 billion long reads—making it one of the largest publicly available long-read datasets to date (Fig.1E, Supplementary Fig.1C,D, Supplementary Table 2). We ensured deep sequencing with at least 23.6 million reads per bulk dataset and 102 million reads per single-cell or single-nucleus dataset, enabling robust gene and isoform detection, accurate quantification, and increased power for downstream analyses, providing a fair basis for cross-platform comparison.

Overall, *LongBench* achieved high-quality data across platforms, with read length and accuracy reflecting inherent technological differences (Fig.1F). The PacBio bulk datasets yielded the largest median read lengths overall (median length 1.53–1.75 kb), followed by ONT direct RNA (dRNA, median length 0.92–1.20 kb) and ONT PCR cDNA datasets (median length 0.59–0.67 kb). However, ONT dRNA produced the longest individual reads (Supplementary Fig.2A). Notably, 10X library preparation yielded similar read lengths between PacBio and ONT (median length 0.74–0.85 kb). These length differences can be linked to library preparation: bulk PacBio Kinnex includes size selection that depletes shorter fragments (0.86–0.9X bead wash after cDNA amplification^(47)^); this step is not present in the single-cell/single-nuclei protocols. Bias against longer transcripts can arise from incomplete reverse transcription (all protocols), premature template-switching (all cDNA protocols) and PCR bias (PCR-based protocols)^(48,49)^. PacBio achieved the highest per-base accuracy and the largest proportion of reads aligned to the genome across the bulk datasets. However, it yielded more intronic reads than the ONT datasets (Fig.2A), likely due to a higher chance of reverse transcriptase priming on internal A-rich regions within pre-mRNA instead of the true poly-A tails (Fig.2B, Supplementary Fig.3). Removal of reads that are likely to have arisen from internal priming (detected using PrimeSpotter, see Methods) sees a substantial reduction in intronic signal (Supplementary Fig. 3C). In contrast, ONT dRNA datasets had the lowest proportion of non-exonic reads (Fig.2A), and importantly, it also uniquely enables reliable detection of RNA modifications (Supplementary Fig. 4). All long-read platforms produced reads that span nearly the full length of transcripts shorter than 1 kb, which represent ~60% of annotated human transcripts. However, the coverage decreased with increasing transcript length, with ONT PCR cDNA showing the most pronounced decline among the three long-read platforms (Fig.2C).

**Fig. 2.**
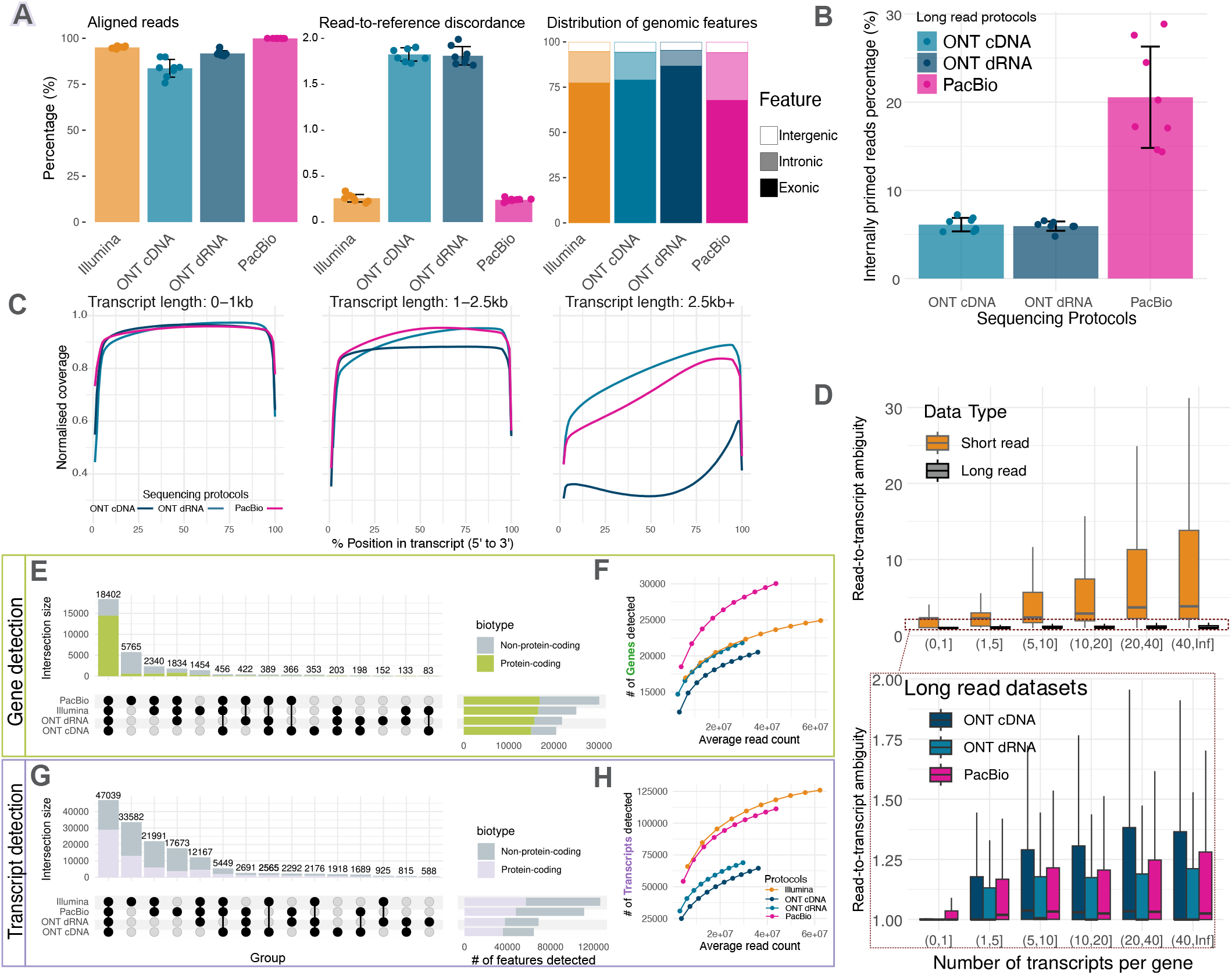
Alignment-based read quality, read-to-gene and read-to-transcript metrics for bulk RNA-seq datasets. **A**. Alignment-based quality summary, including the percentage of aligned reads, read-to-reference discordance (percentage of mismatches, insertions and deletion rates), as well as the distribution of reads across genomic features. **B**. Percentage of internally primed reads in long-read RNA-seq datasets estimated using PrimeSpotter. **C**. Normalised transcript coverage plot. Transcripts were grouped into length bins, and for each group, the normalised coverage (y-axis) is shown across transcript percentiles (x-axis), where 0% corresponds to the 5^*′*^ end and 100% to the 3^*′*^ end of the transcripts. Normalised coverage at a given position percentile was calculated as the number of reads covering that percentile position divided by the total number of reads mapped to the transcripts in the group. **D**. Read-to-transcript ambiguity estimation following the approach of Baldoni et al. ^(50)^. **E, G**. Overlaps of detected genes (E) and transcripts (G) from the GENCODE annotation **F, H**. Rarefaction curve showing the number of genes (F) and transcripts (H) detected when down-sampling the data to account for the effect of different sequencing depths across the datasets.

### Long-read platforms consistently reduce transcript assignment ambiguity

Accurate transcript-level quantification remains a fundamental challenge in RNA-seq analysis, particularly for multi-isoform genes. To evaluate whether long-read platforms consistently reduce transcript assignment ambiguity, we compared the distributions of read-to-transcript ambiguity across protocols. For short reads, ambiguity greatly increased with isoform diversity (Fig.2D), while long reads showed comparatively stable transcript assignment across all protocols, underscoring their advantage in resolving transcript-level expression.

### Differences in detection of expressed genes and transcripts across platforms

We next evaluated how different bulk RNA-seq protocols detect comparable or distinct gene and transcript sets (Fig.2E-H). Among the long-read datasets, PacBio detected the most genes and transcripts. Detected protein-coding features were largely consistent across platforms with Mean Overlap 95.9% and 82.2% at the gene- and transcript-level respectively (see Methods), whereas non-protein-coding features were more platform-specific (Mean Overlap 70.2% and 67.0% at the gene- and transcript-level respectively).

Within long-read datasets, PacBio detected twice as many non-protein-coding features (Fig.2E,G). However, the PacBio reads do not cover significantly more splice junctions with matched depth (Supplementary Fig. 2B), and over 60% of PacBio-specific genes are single-exon genes that overlapped with introns of other genes (Supplementary Fig.5A). Given the higher proportion of intronic reads and propensity for internal priming in PacBio, a possible explanation is that some reads assigned to non-coding genes originate from intronic regions amplified from incompletely spliced overlapping coding transcripts, making false positive detections (Supplementary Fig.5A,B). Filtering out reads likely to have arisen from internal priming substantially reduced the detection of intron-overlapping genes (Supplementary Fig. 6C). In addition, the preferential detection of transcripts and genes with lower GC content by PacBio may have also contributed to PacBio-specific detection (Supplementary Fig.8). While oarfish^(34)^ served as the default long-read quantification tool (see Methods), we note that the potential false positive detection can be reduced computationally by using tools such as IsoQuant^(35)^, which have more stringent read-to-transcript assignment (Supplementary Fig.6C, Methods); however, this can result in reduced overall sensitivity in gene and transcript detections, particularly the short transcripts (Supplementary Fig.6) in PacBio data.

In contrast, ONT cDNA missed many longer genes and transcripts that were detected by all other platforms (71.4% of genes and 84.2% of transcripts specifically missed by ONT cDNA are longer than 1.5kb (Supplementary Fig.5C,D), consistent with the read length distribution observed (Fig.1F) and finding of limited coverage for long transcripts (Fig.2C). Notably, short-read data reported significantly more transcripts than long-read data at matched read depth (Fig.2H), due to the higher ambiguity in isoform assignment (Fig.2D), leading to the assignment of reads to reference isoforms not actually expressed in the sample^(51–53)^.Analysis of the SIRV spikeins (see Methods) highlights this, with more reads assigned to false positive “over-annotated” transcripts (Supplementary Fig.7A) and more false positive transcripts detected (Supplementary Fig.7B) in the Illumina short-read data than in the long-read platforms. Taken together, the different limitations demonstrated by PacBio (intronic reads, internal priming, possible false positive detections or reduced sensitivity with stringent read to transcript assignment), ONT cDNA (poor coverage of longer transcripts, false negative detections), and Illumina (ambiguous isoform assignment) suggest ONT dRNA may provide a more accurate view of transcribed genes and isoforms.

### Large variation in transcript quantification across platforms

We next examined how protocol choice affected transcript quantification. To minimise the influence of differences in gene and transcript detection on quantification and downstream differential analyses, only commonly detected features were retained. We first compared quantification accuracy using spike-in controls with known concentrations (Fig.3A-B, Supplementary Fig.9, Supplementary Information Section 1.1). ONT dRNA consistently showed the highest agreement between observed and expected transcript abundances with the lowest overall normalised root mean square error (NRMSE). Length-dependent platform-specific biases were evident: shorter spike-ins were under-estimated in PacBio, consistent with the longer length distribution noted above and likely influenced by the protocol’s size-selection step, whereas longer ones, like long SIRVs, were underestimated in other datasets, particularly in the ONT cDNA datasets. Emerging hybrid strategies have the potential to improve quantification by leveraging the complementary strengths of long- and short-read sequencing^(54)^. We applied miniQuant in hybrid mode to our data (see Methods) and observed improvements in quantification accuracy in some contexts (Supplementary Fig.10). For example, hybrid quantification of SIRV transcripts combining Illumina and PacBio data achieved a lower mean squared error (4.55) compared to using Illumina or PacBio data alone (5.57 and 15.03, respectively). However, the length biases observed persisted, indicating the need for such methods to explicitly model platform-specific effects.

**Fig. 3.**
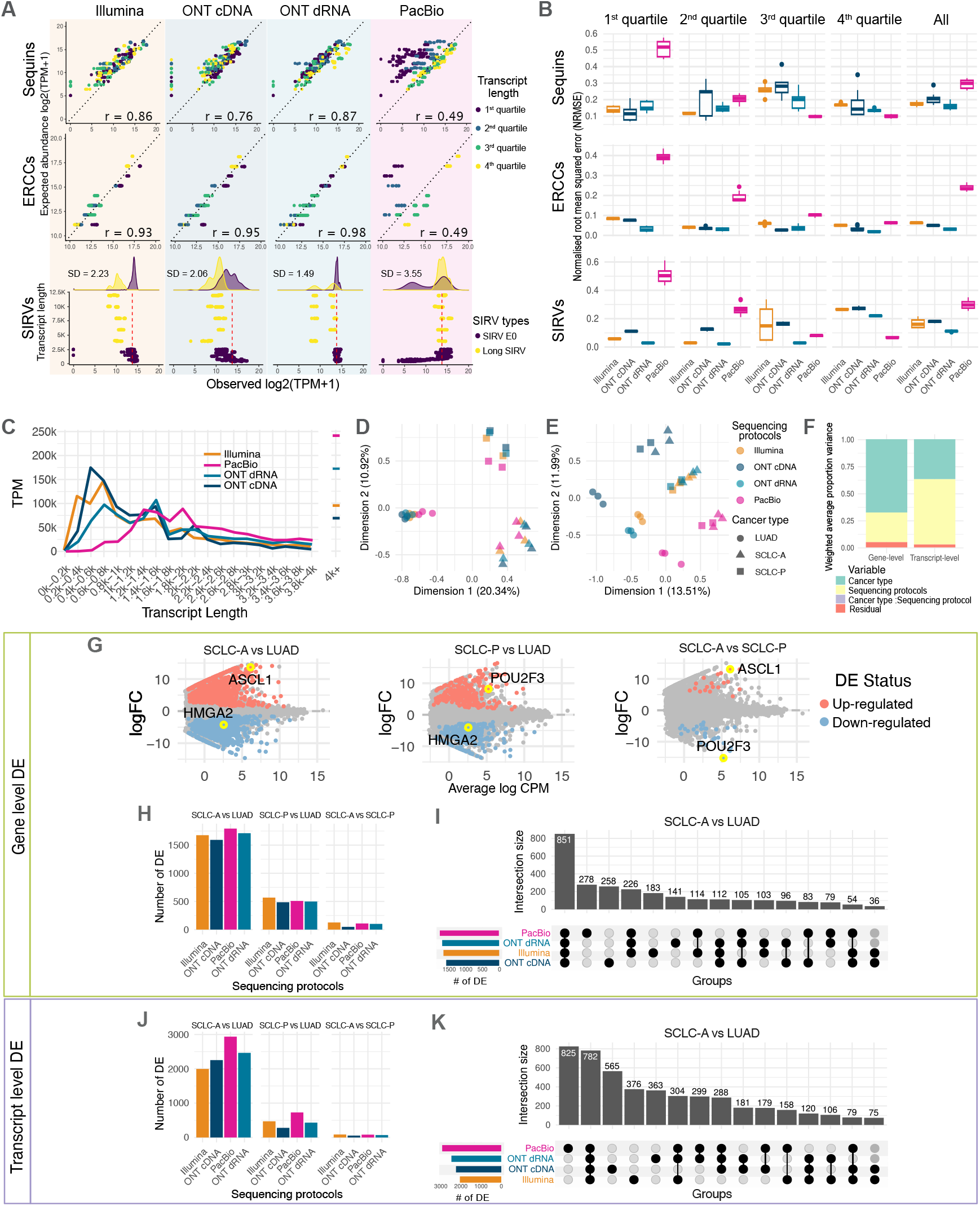
Quantification and differential analysis on bulk RNA-seq datasets. **A**. Comparison of observed log_2_(TPM) of spike-in transcripts to expected abundance. Each dot represents one transcript in one cell line, with all eight cell lines shown in each panel. For Sequins and ERCC, each transcript has a distinct expected abundance (y-axis) and is coloured by transcript length. Pearson correlation (r) is calculated to assess the consistency between observed and expected values. For SIRVs, all transcripts are expected to have equal abundance (indicated by red lines), with transcript length plotted on the y-axis. Standard deviation (SD) and Mean Absolute Error (MAD) are calculated to quantify the biases in quantification. **B**. Normalised root mean squared errors for spike-in transcript quantification in different length quartiles, calculated as: 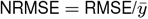, where *RMSE* is the root mean square error and 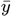 represents the mean expected *log*_2_ (*TPM* + 1) of the spike-in transcripts within each respective quartile. **C**. Count distribution across human transcripts with different lengths. Transcript-level counts were summed across eight cell lines and then converted to TPM. **D, E**. Multidimensional scaling (MDS) plot of samples based on gene-level (D) and transcript-level (E) quantification. **F**. Principal Variance Component Analysis (PVCA) showing the proportion of total variance in gene- and transcript-level quantification explained by each factor and their interactions. **G**. Mean-Average (MA) plot with marker genes of each cancer type highlighted. The plot was generated using the bulk ONT cDNA dataset. The same markers are successfully detected from the other datasets (Supplementary Fig. 14C). **H, J**. Number of gene-level (H) and transcript-level (K) DE identified across three cancer-type comparisons. **I, K**. Upset plots showing the overlap of differentially expressed genes (I) and transcripts (L) identified from each sequencing protocol in the comparison between LUAD and SCLC-A.

A similar pattern was observed for human transcripts, with PacBio undercounting transcripts below 1 kb and ONT cDNA overestimating them relative to other protocols (Fig.3C). Gene-level correlations were generally high, with the strongest concordance observed between ONT dRNA and Illumina datasets (Supplementary Fig.12). In contrast, transcript-level correlations exhibited substantial variability. To assess whether variation reflected underlying biology, we applied MDS and PVCA, which showed that gene-level variation was primarily explained by cancer type, whereas transcript-level variation was dominated by protocol choice (Fig.3D–F). Robustness of this observation was assessed using IsoQuant (see Methods), which yielded consistent conclusions that extended to differential expression and differential transcript usage analyses (Supplementary Figs.10, 11 and 18).

### Bulk RNA-seq differential expression results are consistent at the gene-level but protocol-dependent at the transcript-level

Building on these observations, we next evaluated how platform-specific differences in transcript quantification impact differential expression (DE) analysis. As an initial validation, all protocols accurately recovered the expected fold changes of spike-in transcripts (Supplementary Fig.13), with ONT dRNA and Illumina showing slightly higher accuracy than the others.

With the 18,402 and 47,039 commonly captured human genes and transcripts, we performed DE analysis between each pair of cancer types. As expected from the MDS plots, the most DE genes and transcripts were found between SCLC-A and LUAD (Fig.3H,J), representing distinct cancer types with three samples each. Fewer DE features were seen in SCLC-P vs LUAD, partly due to lower power (only 2 SCLC-P samples), and the least between the SCLC-A and SCLC-P subtypes.

Overlapping DE genes were highly consistent across sequencing protocols (Mean Overlap 65.2%, Fig.3I, Supplementary Fig.14A,B, Supplementary Table 3, Supplementary Fig.18). DE genes detected by all four protocols included well-established cancer-associated genes. For example, HMGA2, known to be highly expressed in non–small cell lung cancer^(55)^, was significantly upregulated in LUAD across all protocols. Key transcription factors ASCL1 and POU3F3, markers for SCLC-A and SCLC-P, were also consistently upregulated in the expected samples (Fig.3G, Supplementary Fig.14C). Among the sequencing protocols, ONT cDNA showed relatively lower sensitivity, missing the highest number of DE features consistently identified by the other three protocols (Fig.3I, Supplementary Fig.14E, F).

The transcript-level DE results were less consistent across protocols (Mean Overlap: 46.9%, Fig. 3J, K, Supplementary Fig. 14G, Supplementary Table 4), with 29.2% of DE transcripts from each protocol being protocol-specific, particularly among lowly expressed transcripts (Supplementary Fig. 14D). The estimated effect sizes from different protocols showed lower consistency across all tested features compared to the gene-level results, consistent with the increased noise at the transcript-level. However, the estimated expression changes for significantly DE transcripts were broadly consistent across protocols, particularly in terms of their direction (Supplementary Fig. 15), with more than 90% of protocol-specific DE transcripts sharing the same direction of effect with the remaining protocols, suggesting that many represent genuine biological signals.

### Transcript usage analysis shows moderate agreement in major isoform detection but divergent differential results

Beyond expression level, isoforms can be analysed in terms of relative isoform usage. We found that the three long-read protocols (ONT cDNA, ONT dRNA, and PacBio) agreed on the major isoform for 67–69% of the multi-isoform genes, whereas short-read results were notably more divergent (Supplementary Fig.16D), likely reflecting the inherent limitations of short-read sequencing in isoform quantification. Moreover, previously described length biases can compromise isoform proportion estimates, especially when isoforms differ substantially in length. When examining differential transcript usage (DTU), the PacBio dataset identified the most DTU events, while ONT cDNA detected the fewest (Supplementary Fig.16E). All four protocols reliably detected known events from the Sequins transcripts (Supplementary Fig. 16A–C). This confirms the technical ability of each method to detect isoform proportion changes in a controlled setting. However, similar to previous studies^(39,56)^, substantial inconsistencies were observed in human genes (Mean Overlap 32.8%, Supplementary Figs.16F,G, 17 and 18), indicating that biological complexity and limited sequencing depth continue to make accurate DTU detection challenging in real samples.

### Consistency of long-read platforms on single-cell and single-nucleus RNA-seq analysis

Single-cell (SC) and single-nucleus (SN) cDNA libraries were prepared from a mixture of all eight cancer cell lines using the 10X Genomics 3’ assay. The same cDNA was then subjected to short-read (Illumina) and long-read (PacBio Kinnex and ONT cDNA) sequencing to compare performance in gene expression analysis tasks. ONT and PacBio protocols assigned comparable proportions of reads to cells (Fig.4A, Supplementary Fig.19). However, a smaller fraction of reads were assigned to nuclei due to elevated ambient RNA introduced during nuclei isolation^(57–60)^. In addition, a doublet rate that was over two times higher was observed in SN data compared to SC data (Fig.4A).

**Fig. 4.**
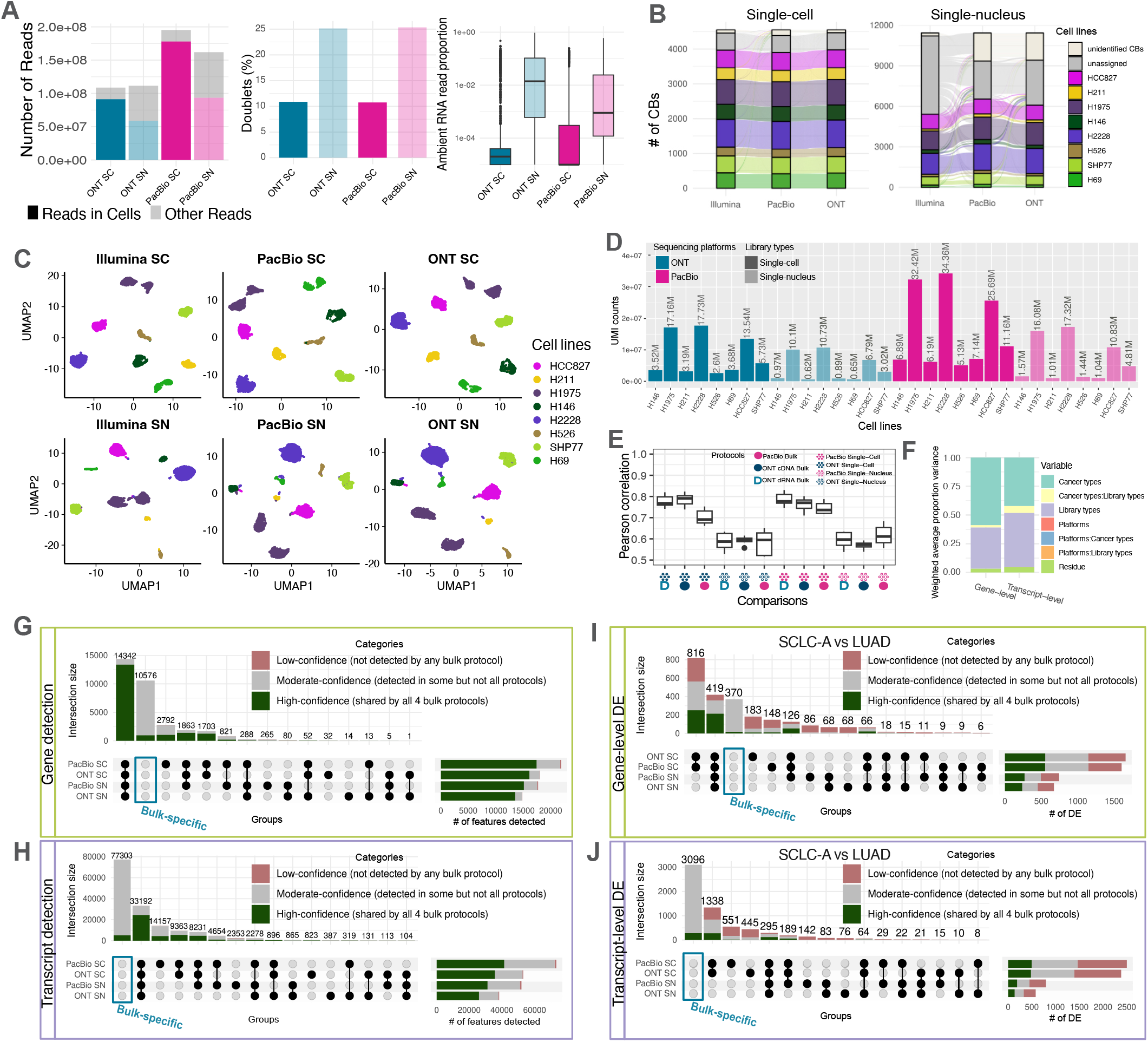
Single-cell and single-nucleus RNA-seq data analysis. **A**. Summary of library quality and read demultiplexing metrics, including the number of reads in cells, doublet rate and proportion of ambient RNA reads. **B**. Genotype-based cell line annotation for single cells and nuclei. Each line connecting the bars represents a matching cell barcode across sequencing platforms. Unassigned cells were excluded from downstream analyses. **C**. UMAP of gene expression profiles from *LongBench* single-cell and single-nucleus datasets, with cells colored by genotype-based cell line labels. **D**. Total UMI counts per cell line among successfully annotated cells. **E**. Pearson correlation of transcript-level quantification between pseudo-bulk and bulk samples was calculated for each cell line, comparing the pseudo-bulk and bulk samples from the same cell line. Only transcripts with non-zero counts across all protocols were included in the calculation. Correlation values were then summarised by the protocol pairs compared, which are indicated on the x-axis. **F**. Principal Variance Component Analysis (PVCA) showing the proportion of total variance in gene- and transcript-level quantification explained by each factor and their interactions across pseudobulk samples. **G, I**. Overlap of detected genes (G) and transcripts (I) in pseudo-bulk samples. Features are classified as low-, moderate-, or high-confidence based on the number of bulk RNA-seq datasets in which they were also detected. **H, J**. Overlap of gene-level (H) and transcript-level (J) DE results in pseudo-bulk samples. DE features are categorised by the number of bulk protocols in which they were also identified as DE.

The identification and SNP-based cell line annotation of single cells and nuclei was consistent across Illumina, PacBio and ONT sequencing platforms (Fig.4B) and validated by expression-based clustering (Fig.4C), with clear separation of all cell lines across all protocols. Notably, H1975 formed two distinct clusters, consistent with previous observations^(61)^. Higher ambient RNA proportion in the SN data again decreased SNP-based annotation success compared to SC (Fig.4B,C). Together with the variation of cell size across the cell lines (the LUAD cells are consistently larger than SCLC cells as confirmed by flow cytometry analysis, Supplementary Fig. 20), cell lines with larger cells/nuclei generated on the order of 10-times more reads, and pseudo-bulk libraries by cell line ranged from 0.65 to 34M reads (Fig.4D). Pseudo-bulk quantification showed moderate to high correlation to matched bulk datasets (Fig.4E), with SC samples correlating better than SN samples. Across 32 pseudo-bulk samples (Fig.4D), library type (SC vs SN) and cancer type explained most variation(94.7% and 89.2% at the gene- and transcript-level), while sequencing platform (PacBio vs ONT) had minimal impact (Fig.4F, Supplementary Fig.19C-F), contrasting with bulk RNA-seq protocols. While individual single cells exhibited sparse feature detection and quantification (Supplementary Fig.19A), pseudo-bulk aggregation recovered most high-confidence features commonly detected in bulk datasets (on average 84.3% and 74.0% at the gene- and transcript-level), although many features seen in bulk remained absent from pseudo-bulk datasets (Fig.4G,H). Down-sampling the bulk data to a similar depth demonstrated the potential for a substantial increase in feature detection with deeper sequencing (Supplementary Fig.19B). Notably, SC and SN datasets detected only a few low-confidence features (<2%), highlighting low levels of spurious feature detection. We then performed gene- and transcript-level differential expression (DE) analysis on high-confidence features commonly detected in SC, SN and bulk datasets (13,376 genes and 24,499 transcripts; Fig.4I, J). DE features were more concordant within the same library type across platforms (Jaccard: 0.39–0.78) than within the same platform across library types (Jaccard: 0.10–0.31), suggesting library type is the primary driver of DE detection. SC datasets identified a comparable number of DE genes as bulk (Fig.3H, 4I and Supplementary Fig.19G) and recovered the majority of high-confidence DE genes (on average 79.1% at both the gene- and transcript-level), whereas the SN datasets showed significantly lower sensitivity (21.7% and 26.6% at the gene and transcript-level respectively). Notably, both SC and SN datasets reported many DE genes and transcripts not detected in any bulk datasets (Fig.4I,J). Given the higher read depth and statistical power in bulk, a substantial portion of these additional SC/SN-specific DE features may represent false positives.

### Variants and allele-specific analyses

RNA-seq also enables the detection of small variants (i.e. SNVs and small INDELs up to 50 nt), gene fusions, and the investigation of allelic differences in gene expression. Using the *LongBench* bulk datasets, we evaluated small variant calling and fusion calling across four sequencing protocols (Fig.5A-E, Supplementary Table 5). PacBio detected the largest number of small variants and gene fusions, predominantly due to higher sequencing yield (i.e., total bases sequenced, Fig. 1F) and greater coverage of intronic regions, identifying more intronic variants (Fig.5B, Supplementary Fig.22). After controlling for sequencing yield (Fig.5D, E), ONT dRNA sequencing demonstrated the highest sensitivity in detecting known mutations and fusion gene pairs, followed by PacBio, while ONT cDNA and Illumina showed lower sensitivity. Notably, long-read datasets demonstrated the potential to detect multi-gene fusion events (≥3 genes, Supplementary Table 5), with a four-gene fusion identified in cell line H69 that all long-read datasets were able to phase across breakpoints (Supplementary Fig.23), reflecting the ability of long reads to resolve more complex fusion architectures that are difficult to reconstruct from short-read data.

**Fig. 5.**
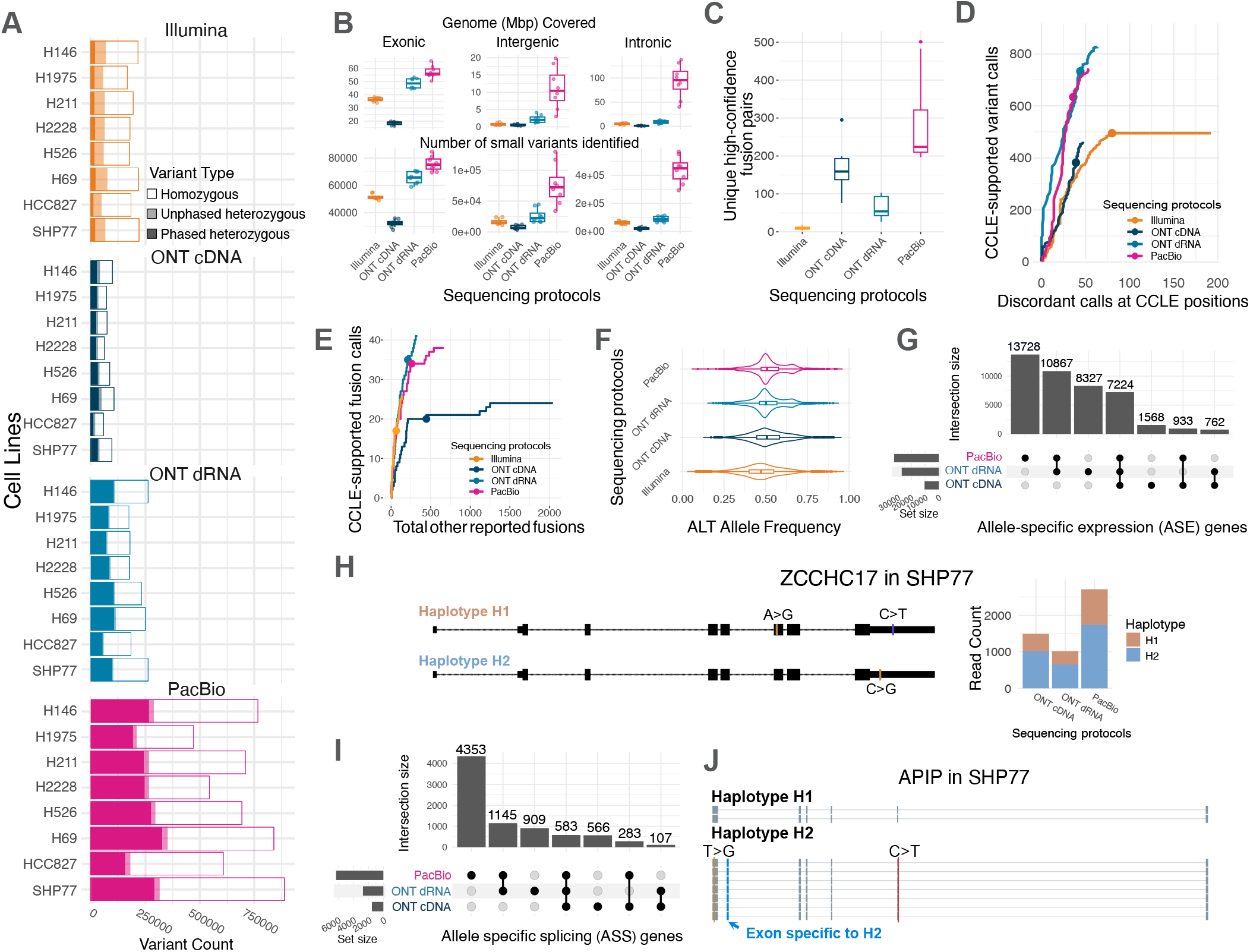
Variant calling and allele-specific analyses: **A**. Number of small variants called by Clair3-RNA (long reads) and Clair3-Illumina (short reads), phased using WhatsHap. **B**. Coverage and number of variants detected across different genomic regions.A minimum read depth of 30 was required for a genomic position to be considered covered. **C**. Number of gene fusions detected by JAFFAL (long reads) and JAFFA (short reads). Each box summarises results across eight cell lines within each protocol. **D**. Detection of known somatic mutations annotated in the Cancer Cell Line Encyclopedia (CCLE). Different numbers of variant calls at CCLE-reported loci were obtained by varying the variant quality (QUAL) threshold from the maximum observed value down to QUAL *≥* 1. The solid dots represent a threshold of QUAL *≥* 10, which was used to report the total number of calls in the panel (A and B). To ensure a fair comparison across sequencing protocols, the input long-read datasets were downsampled to match the total gigabases of the short-read data (see Methods). **E**. Cumulative number of CCLE-supported fusions (y-axis) against other fusion calls (x-axis) across all cell lines, derived from a ranked list of fusion candidates. To ensure a fair comparison across sequencing protocols, the input long-read datasets were downsampled to match the total gigabases of the short-read data (see Methods). Fusion calls from all eight cell lines were aggregated for each sequencing protocol. The solid dots represent a threshold of spanning reads *≥* 3, which was used to report the total number of fusion calls in panel C. **F**. Alternative allele frequency distributions from Clair3-RNA and Clair3-Illumina across datasets. **G**. Allele-specific expression (ASE) genes from different bulk long-read datasets identified using LongcallR. The analysis is performed independently on each cell line, and the sum of the results is presented. A gene detected in multiple cell lines is counted once per cell line. **H**. Example of the ASE gene *ZCCHC17* in SHP77, consistently identified across all long-read protocols; haplotype-distinguishing variants are highlighted. **I**. Overlaps of allele-specific splicing (ASS) genes from different bulk long-read datasets identified using LongcallR. The analysis is performed independently on each cell line, and the sum of the results is presented. A gene detected in multiple cell lines is counted once per cell line. **J**. Example of the ASS gene *APIP* in SHP77, consistently identified across all long-read protocols. An exon is specifically observed in one haplotype.

Beyond variant detection, RNA-seq enables investigation of allelic imbalance analysis, which measures deviations from the expected 50/50 expression ratio at heterozygous loci. This can arise from *cis*-acting regulatory variation, imprinting, or post-transcriptional effects^(62,63)^. Examining alternative allele frequencies, all long-read datasets showed medians near 0.5, indicating balanced expression and unbiased mapping, whereas Illumina displayed a slight reference-allele skew (Fig.5F), and more allelic imbalance events were detected in Illumina at matched depth (Supplementary Fig.21C–E), likely due to reduced mappability of alternative-allele reads. To mitigate the mapping bias, a common issue in short-read data, conventional methods include masking variant positions in the reference genome, making personalised references. However, such approaches may be less critical for long-read sequencing, which spans larger genomic regions and provides a simpler workflow.

While conventional allele-specific analysis focuses on individual heterozygous sites, haplotype-aware analysis links full-length transcripts to specific haplotypes, revealing coor-dinated regulation, allele-specific splicing, and cis-regulatory effects missed by site-level approaches^(64)^. Short-read RNA-seq is limited in haplotype reconstruction, phasing only a small proportion of heterozygous variants (Fig.5A). In contrast, long reads span multiple variants per transcript, enabling phasing of the majority of sites and increasing confidence in variant calls, as spurious errors are less likely to be phased with real variants (Supplementary Fig.21A,B).

Phasing variants with long reads enables the detection of allele-specific isoform usage and improves robustness by aggregating signals across multiple phased variants. Using the long-read datasets, we performed haplotype-resolved allele-specific expression (ASE, Supplementary Table 6) and allele-specific splicing (ASS, Supplementary Table 7) analyses to identify haplotype-dependent differences in gene expression and isoform structure (Fig.5G–J). PacBio detected the highest number of ASE and ASS events, followed by the ONT direct RNA dataset, with many events consistently identified across all three long-read platforms (Fig.5G, H). For example, the ASE gene *ZCCHC17* in SHP77 was detected across all protocols, showing a clear difference in expression levels between haplotypes (Fig.5H), and the *APIP* gene showed haplotype-specific exon usage (Fig.5J). An additional example is provided in Supplementary Fig.21F,G.

## Discussion

*LongBench* builds on the groundwork established by large-scale benchmarking efforts such as LRGASP^(40)^ and SG-NEx^(41)^, while addressing several of their key limitations. These studies were limited to bulk RNA-seq and relied on older sequencing chemistries that do not reflect the performance of current platforms. In contrast, *LongBench* spans bulk, single-cell, and single-nucleus RNA-seq across state-of-the-art long-read technologies. Its experimental design with distinct but biologically related cell lines is explicitly geared towards evaluating DE analysis with realistic and meaningful contrast.

Despite the promise of long-read RNA-seq to deliver quantitative, full-length transcript sequencing, *LongBench* demonstrates that all protocols show specific length and quantitation biases. PacBio Kinnex’s improved throughput enables differential analysis of bulk, SC and SN transcriptomes with highly accurate reads, though cost remains higher than other platforms and transcript lengths are constrained at both ends of the spectrum, particularly the under representation of transcripts below 1 kb with the standard bulk protocol^(42)^. ONT cDNA sequencing is less costly but yields shorter and less accurate reads, with reduced sensitivity for gene, transcript, splice junction detection and DE analysis. However, these differences may mainly arise from differences in manufacturer-recommended library preparation protocols; in single-cell and single-nucleus libraries, where cDNA is prepared uniformly, the choice of sequencer has only subtle effects in these analyses. ONT direct RNA sequencing had the lowest bias of bulk transcriptomic methods, while the accuracy and throughput improvements of the RNA004 chemistry enable confident isoform assignment and SNP analyses to complement its unique ability to detect RNA modifications. However, direct RNA requires relatively large RNA inputs, is slightly more costly than ONT cDNA and has lower accuracy. It also lacks compatibility with single-cell applications and provides limited barcoding options.

Although several previous studies have reported high variability in transcript quantification^(39–41)^, *LongBench* uniquely incorporates realistic biological variability, enabling more representative differential analyses. Consistent with previous observations of transcript-level variability, logFC estimates were generally less consistent across platforms than at the gene-level. Nevertheless, commonly used differential expression testing methods remained relatively robust in detecting true signals. From a practical perspective, DE transcripts identified across platforms may be combined to partially compensate for platform-specific transcript-capture biases. In contrast, overlap in DTU events across platforms was substantially lower, indicating that platform choice itself represents a major source of variation that should be explicitly modelled or stratified by rather than ignored. Consequently, platform-specific DTU hits should be interpreted cautiously and treated as candidate events requiring independent validation. These findings may help guide technology selection during experimental design and inform downstream interpretation of transcript-level analyses.

Hybrid approaches may allow sequencing platforms to complement each other for more accurate transcript quantification. While methods exist for isoform identification^(33,35,65,66)^, few studies have explored combining long and short reads for transcript quantification^(54,67,68)^. Our analyses also suggest the potential for improved quantification through combining long- and short-read sequencing, although the platform-specific biases observed here are likely to persist unless explicitly modelled. As sequencing costs decline, integrating multiple long-read platforms could also become a practical way to overcome individual limitations. However, no established framework yet exists for joint analysis across long-read platforms. Resources like *LongBench* can support further method development to determine how best to combine data and harness the complementary strengths of these technologies.

Variant analysis highlighted another major advantage of long reads: the ability to detect and phase variants directly, enabling allele-specific expression and splicing analyses at high confidence. This provides critical insights into regulatory mechanisms such as imprinting, *cis*-regulatory effects, and isoform-level consequences of genetic variation. Furthermore, it lays the foundation for long-read-based expression QTL and splicing QTL analysis^(69,70)^, offering improved resolution in linking genetic variants to transcript-level regulatory effects. Long-read sequencing also holds particular promise for detecting complex structural variants such as gene fusions, where the extended read length enables resolution of multi-gene fusion events involving three or more partners that are challenging to identify with short reads. At present, most commonly used variant callers are not yet optimised for single-cell RNA-seq. Therefore, in this manuscript, we focused on bulk RNA-seq. However, *LongBench* provides a valuable resource for future method development and validation in that direction.

Our deeply sequenced, multi-platform dataset provides a valuable resource for lung cancer cell lines, particularly at the transcript-level, which is often underrepresented in large-scale resources such as CCLE. By integrating complementary technologies, our design helps mitigate platform-specific biases and reduce false discoveries. In this study, although we focused on benchmarking sequencing technologies using pseudo-bulk analyses, the single-cell long-read data could enable high-resolution studies of expression heterogeneity. Technical noise and sparse coverage at the single-cell level may further limit the reliability of DE analyses^(23,44)^, making the matched bulk datasets a critical reference to control for such artefacts.

While *LongBench* provides a systematic comparison of currently available long-read RNA sequencing platforms, we note that sequencing technologies and analysis tools continue to evolve rapidly. The results presented here reflect current implementations at the time of analysis and may not generalise to other chemistries available on these platforms, such as older chemistries that remain in use or new releases. As protocols and flow cell designs continue to evolve, updated benchmarking will be necessary to reflect the latest platform capabilities. Additionally, our findings reflect the sequencing depths achieved in this study, which represent deeply sequenced libraries by current standards. Nevertheless, specific applications, such as detection and quantification of rare isoforms, may benefit from even greater sequencing levels. By establishing a standardised reference-based benchmarking framework and a modular analysis pipeline, *LongBench* can be readily extended to incorporate newly developed sequencing platforms (such as Roche SBX Sequencing, CycloneSEQ Technology, and future development of ONT and PacBio), thereby remaining adaptable to rapid technological advances in the field. We anticipate that *LongBench* will serve as a foundational dataset for the development, validation, and refinement of RNA-seq analysis tools, including those that integrate multi-platform data, capture transcript diversity more accurately, or advance single-cell and allele-specific transcriptomics.

## Methods

### Cell Culture

Our study included adherent cell lines (H1975, HCC827, H2228) and suspension cell lines (H211, H526, H69, H146, SHP77). Cells were maintained in RPMI 1640 medium supplemented with GlutaMAX™ (Thermo Fisher Scientific), 10% heat-inactivated fetal bovine serum (FBS; Gibco), and 1% penicillin-streptomycin (Thermo Fisher Scientific). Cultures were incubated at 37°C in a humidified atmosphere containing 5% CO_2_. Cells were routinely passaged at 70–80% confluency using standard culture procedures.

### RNA extraction for bulk RNA sequencing

For consistency with the single-cell experiment, adherent cell lines were first detached with TrypLE Express (Gibco). For all cell lines, 1.5–2 M cells were collected for TRIzol-mediated RNA extraction. RNA quantity and quality was assessed with a Nanodrop spectrophotometer, Qubit and TapeStation. The yield was 20–80 µg of RNA per cell line with an RNA Integrity Index (RIN) of 10. Total RNA was supplemented with spike-in controls from Sequins mixtures A and B^(46)^, as well as SIRV Set 4 (Lexogen), which includes Iso Mix E0, ERCC, and Long SIRVs. Sequins mix A was added to RNA from SCLC-P and LUAD cell lines, while mix B was used for SCLC-A. Spike-in preparations followed the manufacturers’ protocols. An estimated mRNA content of 2% in the cell lines was used to calculate the spike-in input, targeting approximately 1% of sequenced reads for Sequins and SIRVs. For Oxford Nanopore direct-RNA libraries sequenced on secondary flow cells, the input amount of Sequins prior to library preparation was increased tenfold due to observed throughput issues specific to Sequins on the ONT platform.

### Single-cell and single-nucleus preparation

Suspension cell lines (H211, H526, H146, H69, SHP-77) were prepared for fluorescence-activated cell sorting (FACS). As these cell lines grow in sheets and aggregates, they were difficult to manually separate into singlets and required sorting for input to 10X Capture. Each cell line was collected from culture, washed twice with PBS, and re-suspended in 500 µL FACS buffer (PBS with 2% FBS). Cells were then passed through a 35 µm strainer cap into FACS tubes and were subsequently stained with 7-amino-actinomycin D for live/dead sorting. Cells were sorted on the BD Aria III to collect 270,000 cells per sample. The remaining 3 adherent lines (H1975, H2228, HCC827) were detached using TrypLE Express (Gibco), washed twice with PBS, and resuspended in FACS buffer. Cell counts were performed in triplicate per cell line on the Countess 3 (Thermo Fisher Scientific) to ensure accuracy, and then 270,000 cells per line were combined into a single mixture. The cell mixture was loaded onto the 10X Genomics Chromium according to the manufacturer’s protocol using the 10X Chromium NEXT GEM single-cell 3’ kit (v3.1) with a target capture of 5,000 cells.

Nuclei isolation was performed using a custom protocol based on the Nuclei EZ Prep kit (Sigma-Aldrich) with the remaining 1.7 M cell mixture. Cells were pelleted at 300 x g for 5 minutes at 4°C, then gently resuspended in 300 µL of cold cell lysis buffer (EZ lysis buffer supplemented with 0.2 U/µL RNaseOUT™). Lysis was carried out on ice for 2.5 minutes to release nuclei, followed by centrifugation at 300 x g for 5 minutes at 4°C. The nuclei pellet was resuspended in 500 µL of cold wash buffer(1X PBS, 1% BSA, 0.2 U/µL RNaseOUT™), incubated on ice for 1 minute, then supplemented with a second 500 µL of wash buffer. Nuclei were gently resuspended by pipetting and pelleted at 200 x g for 5 minutes at 4°C. The pellet was resuspended in 200 µL of wash buffer, topped up to 1 mL, and centrifuged at 130 x g for 5 minutes. Nuclei were finally resuspended in 500 µL of wash buffer for quality assessment via DAPI staining and microscopy using Countess 3 system. The suspension was filtered twice using FlowMi 40 µM filter tips (Sigma-Aldrich) to remove any present clumps and re-counted prior to 10X capture. The nuclei mixture was loaded onto the 10X Genomics Chromium following the same protocol as described for single-cell capture, with a target capture of 5,000 nuclei.

### Library preparation and RNA sequencing

#### Bulk RNA sequencing libraries

Illumina libraries we constructed using the Illumina Truseq v2 Stranded kit with polyA selection. Superscript III was used instead of SuperScript II, and the reverse transcription incubation step was accordingly adjusted to 50°C. The pool of eight indexed libraries was sequenced on one NextSeq2000 Mid-Output flow cell (400M 100PE reads). Nanopore direct-RNA sequencing libraries were constructed using the ONT SQK-RNA004 kit, with the following modifications: input was increased to 2 µg total RNA, we omitted the RNA CS (calibrating strand), and used the NEB Induro Reverse Transcriptase rather than the SuperScript III. Four libraries were prepared for each cell line, and sequenced over two PromethION RNA flow cells (FLO-PRO004RA), including one wash (custom wash mix based on the ONT SQK-WSH004 kit of 2 µL WMX, 1 µL RNase Cocktail, 1 µL RNase H, 396 µL DIL) and reload per flow cell (16 flow cells total). Library preparation and sequencing were performed in two runs, each comprising preparation of two libraries and sequencing on one flow cell per cell line. Nanopore PCR-cDNA libraries were constructed using the ONT SQK-PCB114.24 kit, with the following modifications: reverse transcription was performed for 90 minutes at 42°C, followed by 14 cycles of PCR with an extension time of 4 minutes. Barcoded samples were pooled and sequenced on three Promethion R10.4.1 flow cells (FLO-PRO114M, 3 flow cells total). PacBio Bulk Kinnex libraries were prepared with the Iso-Seq express 2.0 kit and Kinnex PCR 8-fold kit at the Australian Genomics Research Facility (AGRF). Individual libraries were sequenced on their own PacBio Revio flow cell (non-SPRQ, 8 flow cells total).

#### Single-cell RNA sequencing libraries

From the cDNA generated by the 10X Genomics single-cell 3’ kit (v3.1, 11 cycles of PCR), libraries were prepared for Illumina, Nanopore and PacBio sequencing. Illumina scRNA-seq libraries were prepared with the 10X Library Construction Kit (PN-1000190) and sequenced on one lane of a NextSeq2000 High-Output flow cell (1.8B 100PE reads). The nanopore library was prepared from 10 ng of input cDNA, following the ONT single-cell protocol including the artifact removal step (v9198_v114_revE_06Dec2023). 125 ng of the post-pulldown PCR product were taken to adapter ligation with the ONT SQK-LSK114 kit. The final wash was performed with a 0.5X bead ratio (adjusted from 0.4X) and the short fragment buffer. 50 ng of the final library were loaded onto one Promethion R10.4.1 flow cell (FLO-PRO114M, 1 flow cell total). The PacBio single-cell Kinnex library was prepared with the Kinnex single-cell 12-plex concatenation kit according to the manufacturer’s instructions, which also include an artifact removal step. The final library was loaded onto two PacBio Revio flow cells (2 flow cells total).

#### Single-nuclei RNA sequencing libraries

From the cDNA generated by the 10X Genomics single-nuclei (12 cycles of PCR), libraries were prepared for Illumina, Nanopore and PacBio sequencing. Illumina snRNA-seq libraries were prepared following the same protocol as the single-cell library and sequenced on one lane of a High-Output flowcell (1.8B 100PE reads). A nanopore single-nuclei library was prepared following the same protocol as the single-cell library, and sequenced on one PromethION R10.4.1 flow cell (FLO-PRO114M, 1 flow cell total). A PacBio single-nuclei Kinnex library was prepared identically to the single-cell protocol. The final library was loaded onto two PacBio Revio flow cells (non-SPRQ, 2 flow cells total).

#### Data pre-processing and quality control

The ONT datasets were basecalled with Dorado using *rna004_130bps_sup@v5*.*1*.*0* for direct RNA and *dna_r10*.*4*.*1_e8*.*2_400bps_sup@v5*.*0*.*0* for the cDNA datasets; reads with mean quality scores *<*10 were discarded. The ONT dRNA data from the two runs for each cell line were pooled for analysis unless otherwise specified, after confirming high concordance and minimal batch effects (Supplementary Table 8). The read length and quality distribution were summarised using Nanoplot. The reads were mapped to the reference genome GRCh38 and GENCODE v44 transcriptome using minimap2 ^(71)^ (v2.28). The resulting BAM files of genome alignment were subsampled to 3 million reads per sample to estimate the alignment rate, alignment-based error and the distribution of the genomic feature using AlignQC in Fig.2A.

#### Internal priming detection

To identify internal priming artifacts in RNA-seq data, we developed a custom tool, PrimeSpotter (https://github.com/youyupei/PrimeSpotter). Briefly, the genome was scanned using a sliding window of 10 nucleotides to identify A-rich regions containing ≥8 “A”s on the forward strand (or ≥8 “T”s on the reverse strand). Overlapping or adjacent windows were merged into contiguous regions and extended by 10 nucleotides upstream and downstream to account for alignment variability. A-rich regions overlapping annotated transcript termination sites were excluded to avoid false positives. A read was classified as internally primed if its 3’ end terminated within an identified A-rich region; reads starting within a T-rich region were similarly flagged, as T-rich regions on the forward strand correspond to A-rich regions on the reverse strand and represent the equivalent artifact for reverse-strand genes.

### Downsampling analysis

Both read-count-based and sequencing-yield-based downsampling analyses were included to assess the effect of sequencing depth on cross-platform comparisons, while all primary analyses used the full datasets unless otherwise specified. Quantification-based analyses, including gene and transcript detection, quantification, and differential analysis, included read-count-based downsampling analysis, where each read contributes equally regardless of read length, and datasets were downsampled to the same number of aligned reads. In contrast, analyses more directly related to transcript coverage, such as small variant, fusion, included sequencing-yield-based downsampling, as longer reads contain more sequence information. For the sequencing-yield-based downsampling, ONT cDNA, ONT dRNA, and PacBio datasets were downsampled to 16.9M, 8.5M, and 6.9M reads per cell line, respectively, resulting in total sequencing yields of 102.1–102.6 gigabases across the eight cell lines, comparable to the Illumina dataset (102.5 gigabases).

### Quantifying feature overlap across platforms

We defined “Mean Overlap” to quantify the consistency of detected features or DE features across the four sequencing platforms. For any given platform, we calculated the proportion of its features also identified by at least two of the three remaining platforms, averaging these proportions across all four platforms. For the differential expression (DE) analysis, this metric was further averaged across the three pairwise comparisons.

### Gene and transcript quantification

Transcript-level quantification was performed using salmon^(72)^ (v1.10.2) for short-read datasets and oarfish^(34)^ (v0.6.2) for long-read datasets, using the GENCODE v44 transcriptome supplemented with spike-in transcripts for bulk datasets (not included in pseudo-bulk quantification). Before quantification, read-to-transcript alignment is essential; Salmon incorporates an in-built alignment process, while minimap2 ^(71)^ (v2.28) was used for long-read alignment with oarfish as mentioned above. Both salmon and oarfish can compute bootstrapped abundance estimates. SIRV “over-annotation” analysis (Supplementary Fig. 7) was performed using the same quantification approach described above, but with the official SIRV over-annotated GTF provided by Lexogen. This GTF contains 69 genuine SIRV transcripts (excluding Long SIRVs) alongside 31 artificially added non-existent isoforms, enabling assessment of quantification robustness in the presence of spurious transcript annotations. Gene-level quantification was obtained by aggregating transcript-level quantification using tximport^(73)^ (v1.34.0). While oarfish served as the default quantification tool, we additionally used IsoQuant^(35)^ (v3.10.0) as an independent validation of our observations, and miniQuant^(54)^ (v1.4.1) to explore potential improvements from mitigating protocol-specific quantification bias by hybridising long- and short-read data. IsoQuant was run using the GRCh38 reference genome and the GENCODE v44 transcript annotation, with the `--no_model_construction` option to restrict analysis to annotated transcripts only. MiniQuant was run using long- and short-read FASTQ files for each cell line, together with the same reference files used for oarfish and salmon.

### Read-to-transcript ambiguity

During quantification using salmon and oarfish, 50 bootstrap samples were generated per run to assess the technical variance introduced during the quantification step. Following the approach of Baldoni et al. ^(50)^, we used transcript-level bootstrap counts to estimate the extra-Poisson overdispersion parameters, which capture the uncertainty arising from the probabilistic assignment of reads to transcripts. In line with Baldoni et al. ^(50)^, we refer to this overdispersion as read-to-transcript ambiguity throughout the manuscript. The method is described in detail in their work and is implemented in the edgeR package.

### Gene and transcript detection

Gene and transcript detection analyses were based on whether a feature had sufficient counts from the quantification step. Specifically, we used the *filterByExpr* function^(74)^ from edgeR^(75)^, which retained features with counts-per-million (CPM) above a threshold determined from a minimum count and the median library size, and required that this CPM threshold be met in at least two of the eight cell lines in our case. We set the minimum count to 10 for genes and 5 for transcripts. The *filterByExpr* function is typically used to define features suitable for differential expression analysis. Although *filterByExpr* is typically used to define features suitable for differential expression analysis, we applied it here to define “detected features”, ensuring that the criteria for feature presence in the detection analysis are directly comparable to those used in DE analysis.

### Analysis of spike-in transcripts

The spike-in transcripts were quantified together with human transcripts in bulk datasets. SIRV Set 4 includes ERCC transcripts, which are present at half the molar concentration of the SIRV Set 4. However, we observed that the different types of spike-ins are captured with varying efficiency, due to differences in their design and synthesis. Especially, both ONT PCR cDNA and dRNA protocols did not efficiently capture Sequins and ERCC transcripts (Supplementary Fig. 9). To avoid introducing this artificial bias in the comparison, we analysed ERCC transcripts separately from the other SIRV transcripts. Unless otherwise specified, “SIRV” refers only to the standard SIRV mix (i.e., SIRV Mix 0) and the long SIRV transcripts, which were added at equal molar concentrations.

### Filtering and normalisation for quantification comparison

Because the differences in feature detection could obscure meaningful quantitative differences, we restricted the quantification comparisons to the genes and transcripts detected by all platforms. The estimated counts were then normalised to “Transcript per million (TPM)” and the TPM for short-read data were adjusted by the effective length of each transcript. Specifically:

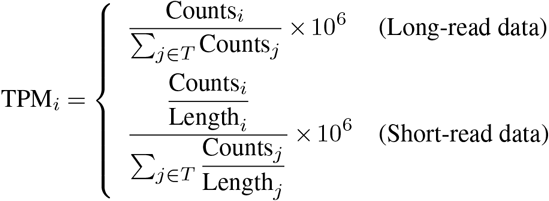

where *T* denotes the set of all transcripts, Counts_*i*_ is the count for transcript *i*, and Length *j* is the effective length of transcript *j*. Note that this TPM normalisation is used only for quantification comparison across sequencing protocols and was performed separately for human transcripts, Sequins, ERCC, and SIRVs. For differential expression analysis across cancer types, raw gene or transcript-level counts were normalised using the strategy below.

#### Differential Expression Analysis

Similar to the quantification comparison, we tested the genes and transcripts detected across all platforms for differential expression (DE). The gene and transcript counts were normalised using trimmed mean of M values (TMM)^(76)^. The transcript counts were further scaled by the overdispersion following the workflow suggested in Baldoni et al. ^(50)^. We used edgeR^(75)^ quasi-likelihood (QL) pipeline to perform gene- and transcript-level DE testing between the 3 groups LUAD, SCLC-A and SCLC-P.

#### Different Transcript Usage Analysis

Differential transcript usage (DTU) analysis was performed using DTUrtle ^(77)^ (v1.0.3), which implements the DRIMSeq^(78)^ framework. Transcript-level counts were generated using tximport (v1.34.0) using raw estimated counts uniformly across all platforms. Per-platform expression-based filtering was applied per pairwise comparison using DTUrtle’s *filtering_strategy = “bulk”* (*min_gene_expr = 5, min_feature_prop = 0*.*05*, thresholds scaled to 50% of the smallest group size, *run_gene_twice = TRUE*). The filtered transcript lists were then intersected across Illumina, ONT cDNA, ONT dRNA, and PacBio per pairwise comparison, so that DTU testing used the same feature set for every protocol across all pairwise Sequins and human comparisons. Significant DTU events at a gene- and transcript-level were identified by applying DTUrtle’s *posthoc_and_stager* function (*ofdr=0*.*05, posthoc=0*.*1*) to the DRIMSeq fitted models for each pairwise comparison. Gene-level significance reflects an overall shift in isoform proportions between conditions, whilst transcript-level significance is determined via a secondary hypothesis test within each significant gene to identify the specific isoforms driving the change; not all significant genes will yield significant transcripts^(77)^.

#### Single-cell and single-nucleus RNA-seq data processing

The short-read single-cell (SC) and single-nucleus (SN) datasets were processed using Cell Ranger (v8.0.1) with GENCODE v44 comprehensive annotations and the GRCh38 primary assembly reference genome. Long-read SC and SN reads were first demultiplexed using BLAZE (v2.5), and the resulting FASTQ files were processed with the FLAMES (v2.0.1) single-cell pipeline to obtain gene-level quantification. Low-quality cells in each SC and SN dataset were filtered out by applying a median absolute deviation (MAD) threshold of 3 on the log-transformed number of detected features (nFeature) and total counts (nCount) to remove extreme outliers. Additionally, cells with mitochondrial gene percentages exceeding 3 MADs above the median were excluded. Cells failing any of these criteria were removed to ensure data quality. After filtering, cells and nuclei commonly detected across Illumina, PacBio, and ONT datasets were retained. Those successfully assigned to a cell line were pooled by cell line within each SC or SN dataset to create pseudobulk FASTQ files. These pseudobulk files were processed identically to bulk datasets to obtain gene- and transcript-level quantification and to perform differential expression analysis.

#### Cell line annotation and detection of doublets and ambient RNA in single-cell and single-nucleus data

Cells and nuclei were annotated based on genotypes at common human SNPs from the 1000 Genomes Project^(79)^. To establish genotypes for the eight cell lines, we first ran CellSNP-lite ^(80)^ on the Illumina bulk RNA-seq data. For each single-cell and single-nucleus dataset, we then ran CellSNP-lite separately to genotype each barcode. Both the bulk and single-cell VCF files were provided to Vireo^(81)^, which assigned each cell or nucleus to the most likely cell line based on its SNP profile. Cell line annotations were obtained independently from the Illumina, ONT, and PacBio datasets. We then merged these annotations by assigning each cell or nucleus to the cell line reported by the majority of platforms. Cells with an equal number of conflicting assignments were discarded from the final merged annotation. Unannotated calls were ignored in the comparison and did not count as disagreement. In each Vireo run, doublets were reported by default and ambient RNA estimation was enabled via the `--callAmbientRNAs` option.

#### Variant calling and allele-specific analyses

Small variant (SNVs and small INDELs up to 50 nt) calling was performed using tools from the Clair3 series^(82,83)^ (v1.0.10), with technology-specific presets. Clair3-Illumina was used with the preset `ilmn` for short-read RNA-seq, and Clair3-RNA^(83)^ was used for the bulk long-read datasets with the presets `hifi_mas_minimap2`, `ont_r10_dorado_cdna`, and `ont_dorado_drna004`. These calls were compared with somatic mutation data for each cell line obtained from the Cancer Cell Line Encyclopedia (CCLE) using BCFtools^(84)^ (v1.21). Next, phasing of heterozygous variants was performed using WhatsHap^(85)^ (v2.8.0). Fusion detection was performed using JAFFA (v2.5, direct mode) for Illumina short-read data and JAFFAL (v2.5) for long-read datasets, with its pre-built reference (GENCODE v49). Known structural variants for the corresponding cell lines were obtained from the CCLE and used for comparison and validation. Finally, allele-specific expression (ASE) and allele-specific splicing (ASS) events were identified with the LongcallR pipeline v1.11.0 ^(64)^, which is tailored for SNP detection, phasing and allele-specific analyses based on long-read RNA-seq data.

#### Software choice for benchmarking analysis

In this study, we focused on comparing different sequencing protocols. To ensure that observed differences arise from the data rather than from analysis variability, we standardised the computational workflow as much as possible. We aimed to select well-established and tested tools that enabled fair and consistent comparisons between short- and long-read sequencing data wherever possible. For transcript quantification, we used salmon and oarfish—tools developed by the same group with similar models. Similarly, we applied Clair3-Illumina and Clair3-RNA for small variant calling, and JAFFA and JAFFAL for fusion calling. Single-cell and single-nucleus datasets were aggregated into pseudobulk profiles and processed through the same pipeline as bulk data (Fig.1D). To minimise tool-specific bias, we also validated the robustness of our key observations using independent analytical approaches: IsoQuant was used as alternative quantification tool to confirm conclusions related to gene and transcript detection, quantification, and differential analysis. Cell line annotations were further assessed through cross-tool comparisons to ensure robust cell-type assignment. The mutation analysis results were evaluated against known ground-truth variants, supporting the validity of the detected signals. However, the limited availability of tools for allele-specific analysis, together with the lack of suitable benchmarking datasets with established ground truth, remains an important limitation. Manual inspection of selected loci through visual examination of read alignments further suggested that some detected signals were plausible, although the overall false-positive rate remains unclear.

## Data Availability

The *LongBench* data are available at https://github.com/mritchielab/LongBench.io. Sequencing data are available from Gene Expression Omnibus under accession number GSE303762.

## Code Availability

The code for the analysis can be found at https://github.com/youyupei/LongBench_workflow. The custom tool that identifies internally primed reads is available at https://github.com/youyupei/PrimeSpotter.

## Author Contributions

Y.Y., A.N.S., J.L., M.D. and C.W. conducted data analysis, generated the figures and wrote the manuscript with input from all authors. J.W.T., K.Z., S.S., C.P., R.G., M.C., J.G. and Y.D.J.P. conducted data analysis. J.L. and Q.G. generated benchmarking data. J.N. conducted the flow cytometry analysis. B.D., I.C. and N.M.D. supervised research. M.L., J.N. and K.D.S. designed the study design, provided cell lines and culture protocols and interpreted results. M.B.C., Q.G. and M.E.R. obtained research funding, designed the study and supervised the research. All authors read and approved the final manuscript.

## ACKNOWLEDGEMENTS

We are grateful to the WEHI Genomics Lab (Casey Anttila, Johannes Wichmann and Daniela Zalcenstein) for processing the single-cell mixtures using the 10x platform, the Australian Genome Research Facility (Cath Moore, Peter Lau, David Hawkes, Angus Li and Trent Peters) for performing PacBio long-read sequencing, Ricardo De Paoli-Iseppi for sharing direct RNA sequencing advice, Paul Gooding (Millennium Science), James Miller and Elizabeth Tseng (PacBio) for advice on sequencing protocols and data analysis and Qingwang Chen for discussions on data analysis. We thank AWS for providing access to the *LongBench* data through the AWS Open Data Sponsorship Program.

