## Supplementary Figures and Information for "Benchmarking long-read RNA-sequencing technologies with *LongBench:* a cross-platform reference dataset profiling cancer cell lines with bulk and single-cell approaches"

### Supplementary Figures and Information from "Benchmarking long-read RNA-sequencing technologies with *LongBench*"

#### Supplementary Figures

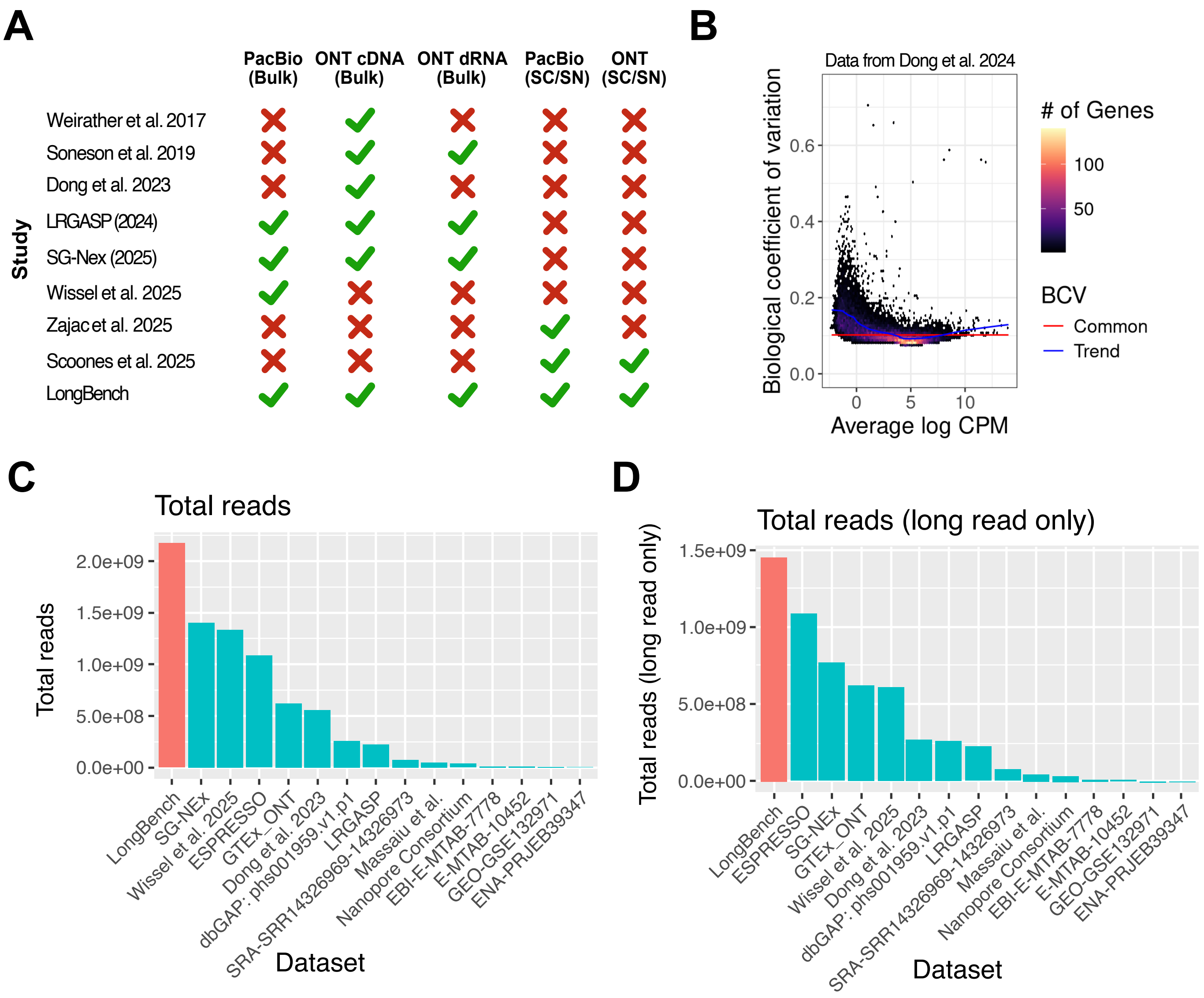

**Supplementary Fig.** 1: **Comparison of long-read RNA-Seq benchmarking studies: A.** Summary of long-read RNA-seq protocols evaluated in existing benchmarking studies[1–8], with the *LongBench* study included for reference. **B.** Example of Biological Coefficient of Variation (BCV) from technical replicates where the same cell line was grown multiple times and sequenced. The data were from Dong et al.[3]. **C-D.** Comparison of total (C) and long-read counts (D) with previously published long-read studies. Most of the values were sourced from Chen et al.[5], with additional data from Wissel et al.[6].

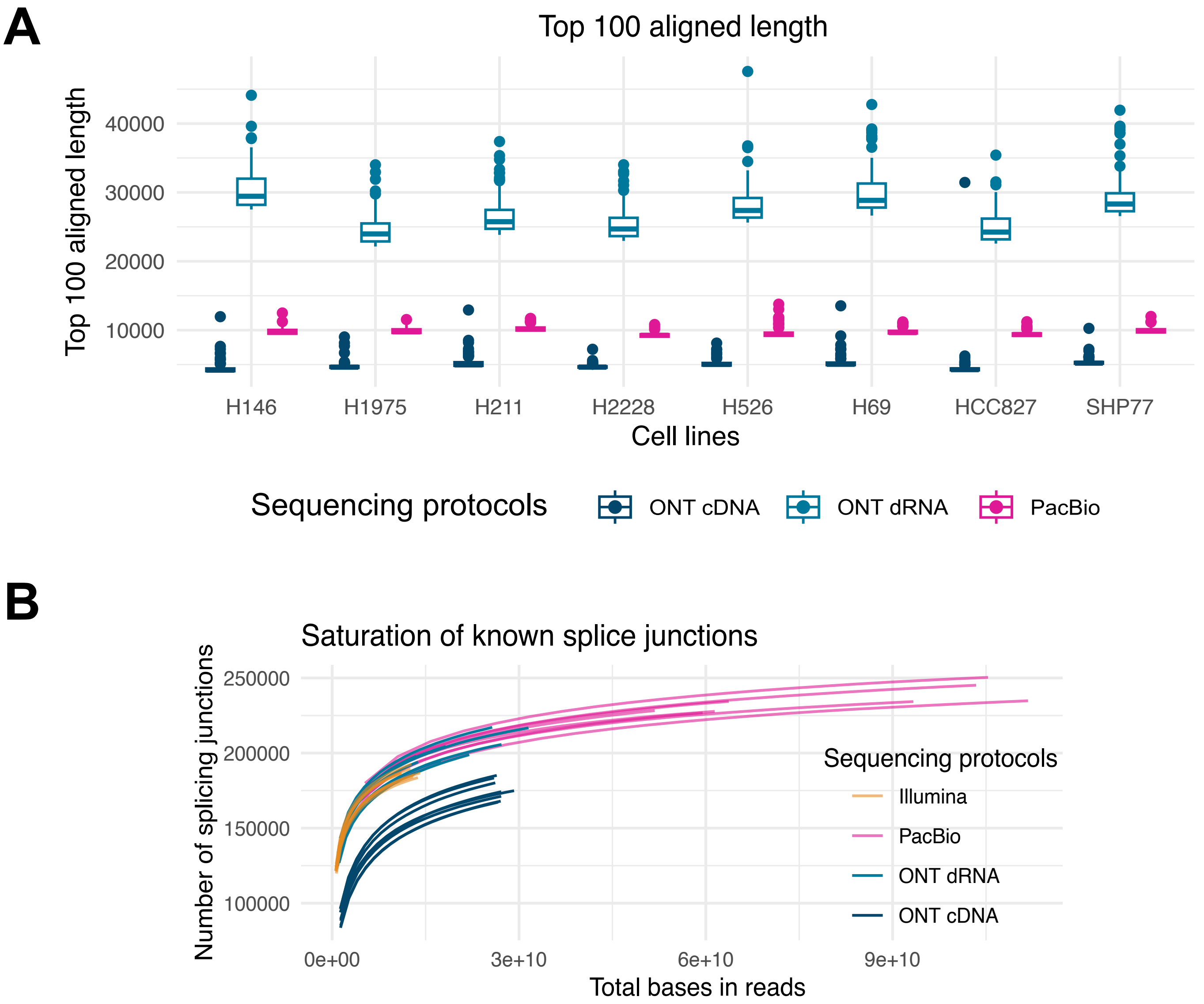

**Supplementary Fig.** 2: **Bulk RNA-seq data quality (extended): A.** Distribution of the 100 longest aligned read lengths per cell line, following mapping to the human genome. **B.** Saturation curve generated using RSeQC[(9)]. The y-axis shows the number of annotated splice junctions detected when downsampling the number of reads. The x-axis represents the number of total sequenced bases. There are 8 curves per sequencing platform representing 8 different cell lines.

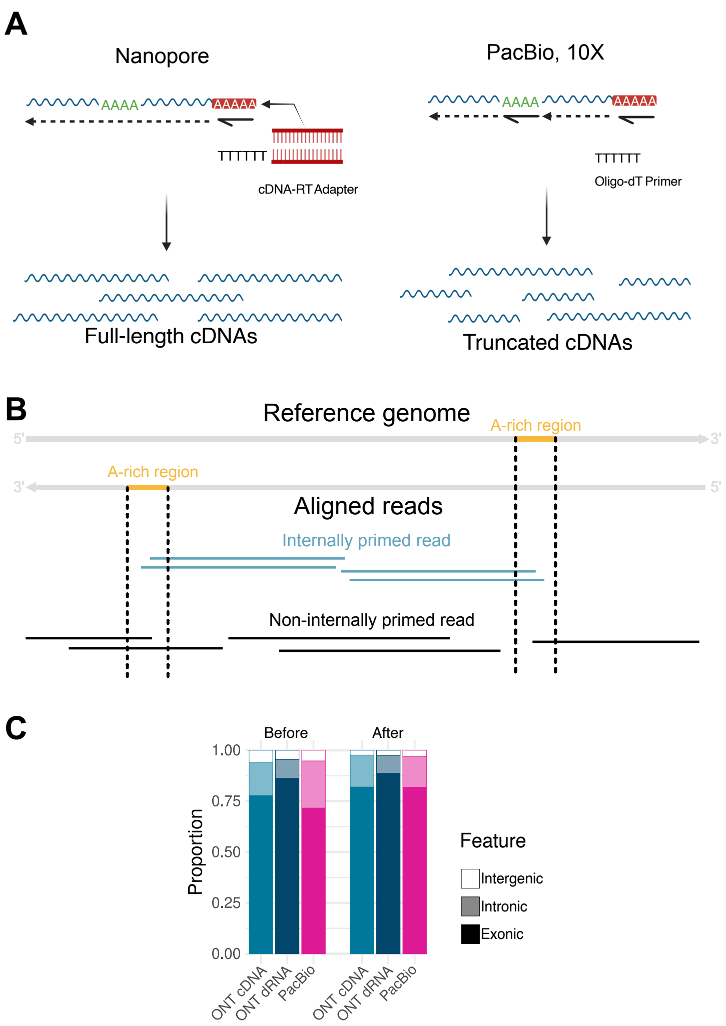

**Supplementary Fig.** 3: **Reverse transcription strategies and internal priming detection: A.** The Oxford Nanopore cDNA-RT Adapter contains a poly-T overhang, allowing reverse transcription (RT) from the very end of mRNA polyA-tails during cDNA synthesis. PacBio Kinnex or 10x Chromium 3' GEX uses oligo-dT for RT, which can prime any polyA signal, causing internal priming and increased prevalence of truncated cDNA. **B.** Overview of the algorithm to classify internally primed reads. Aligned long reads indicating a transcript termination site within an A-rich genomic region were classified as internally primed, since true polyadenylated tails are not encoded in the genome. Note that, for each A-rich region, we only consider reads that align to the 5' side of the region on the same strand. The detailed implementation can be found at `https://github.com/youyupei/PrimeSpotter`. **C.** The proportion of intergenic, intronic and exonic long reads before and after removing internally primed reads.

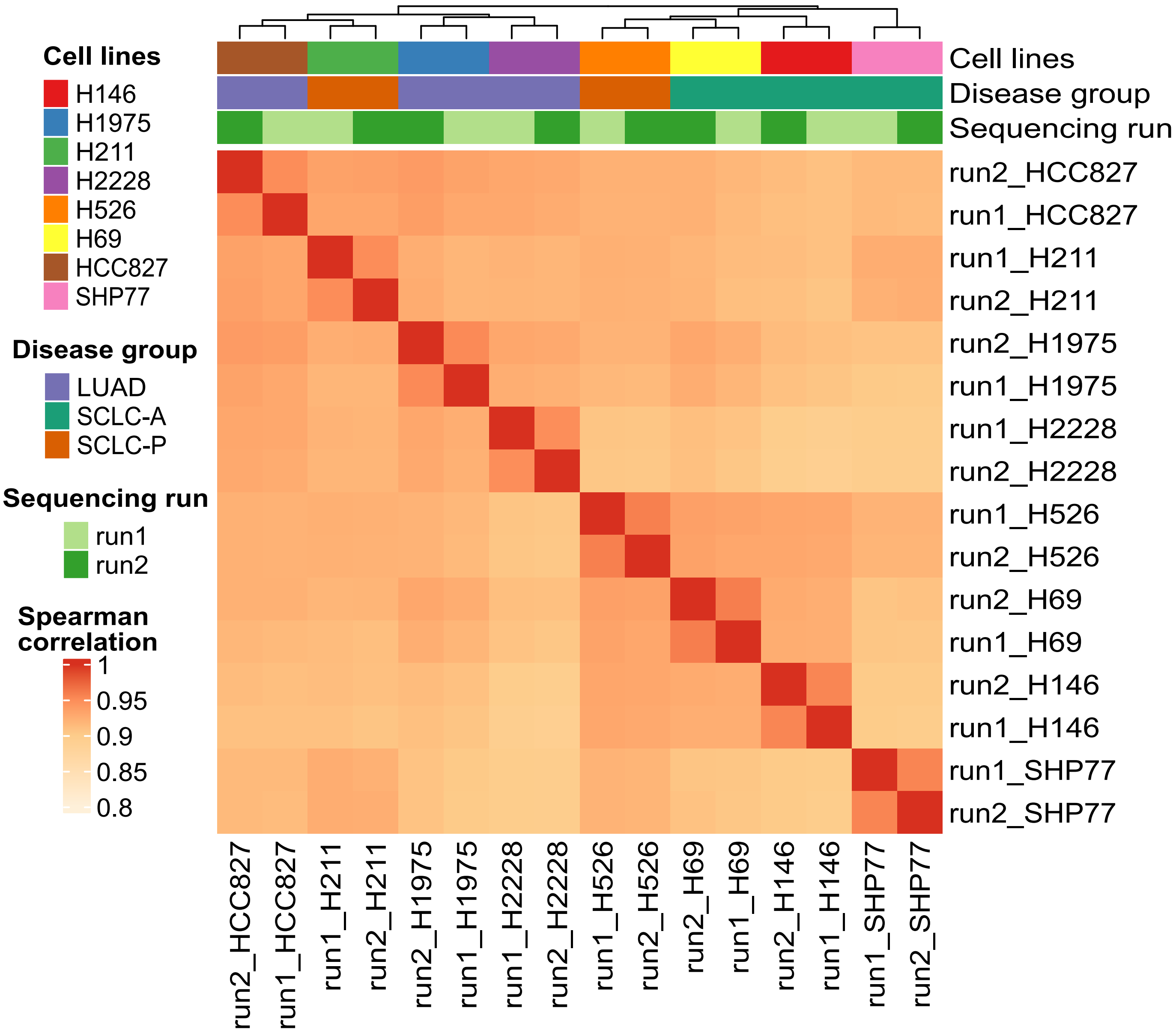

**Supplementary Fig. 4: Detection of RNA modification using ONT directRNA sequencing:** Spearman correlation of modification proportions across 221,809 DRACH loci across all chromosomes (after filtering such that each locus had >20 coverage in each sample and showed >20% modification in at least 2 samples). Two independent sequencing runs per cell line were basecalled with Dorado (rna004_130bps_sup@v5.1.0 model). The consistently high correlations demonstrate that modification profiles are highly reproducible between runs.

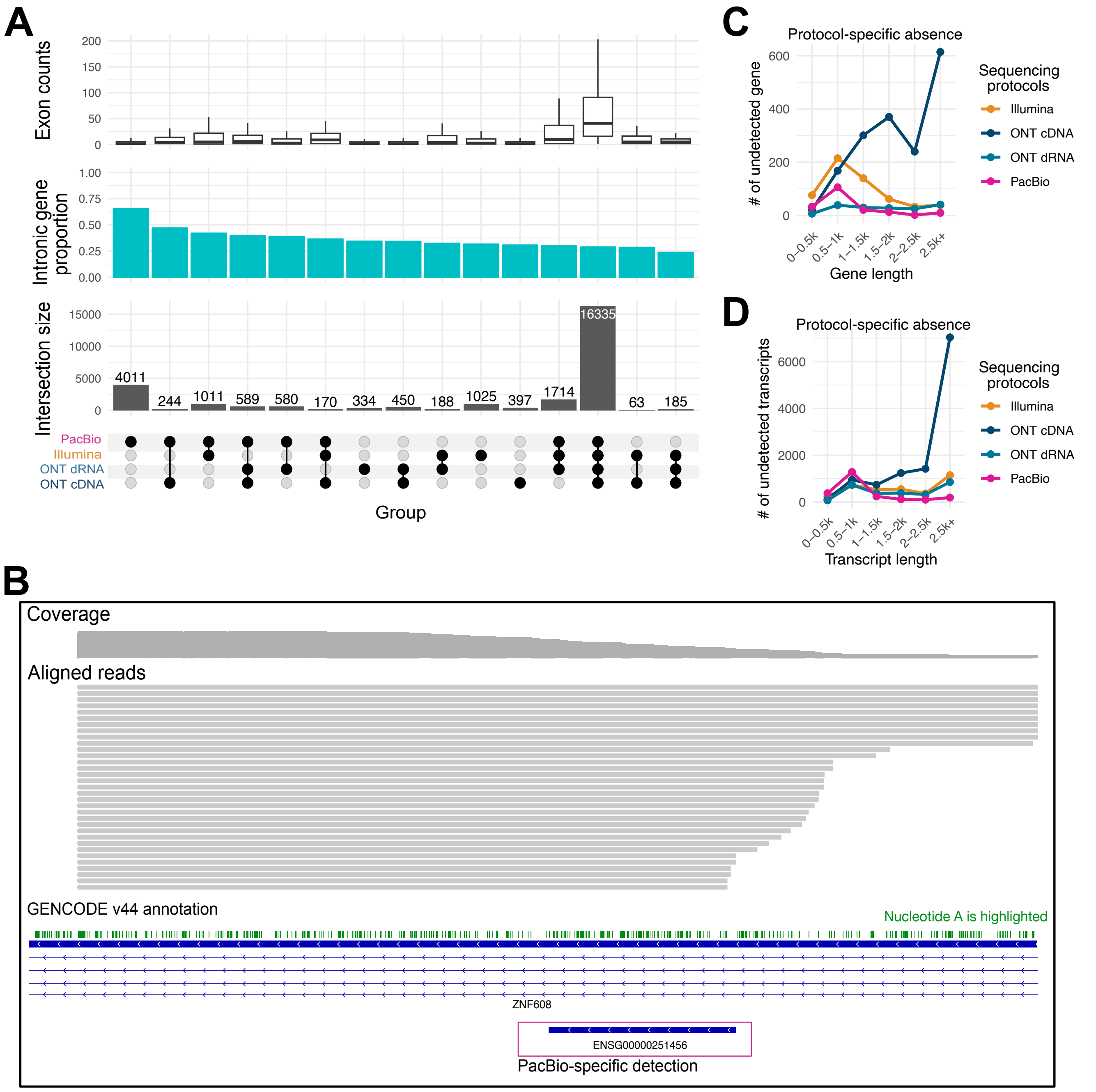

**Supplementary Fig.** 5: **Potential false positive and false negative feature detections:** Each bulk RNA-seq datasets were downsampled to 20 million reads per cell line to match the read depth. **A:** Overlaps of detected genes. The bottom panel of A shows the overlap of genes detected across four bulk RNA-seq datasets. The panels above display various gene features, including: (1) intronic gene proportion — the fraction of genes whose exons overlap with introns of other genes; and (2) exon count — the number of annotated exons per gene. **B:** An example of a gene specifically identified in the PacBio dataset. Aligned PacBio reads are visualised in Integrative Genomics Viewer (IGV) at chr5:124,733,070–124,736,069 for the H146 sample. ENSG00000251456 was detected only in PacBio. The aligned reads originate from an A-rich region within an intron of ZNF608, likely resulting from internal priming, and extend into ENSG00000251456. **C-D:** Length distribution of protocol-specific undetected genes (C) and transcripts (D) that are detected in all other protocols. Gene length was calculated as the average transcript length per gene.

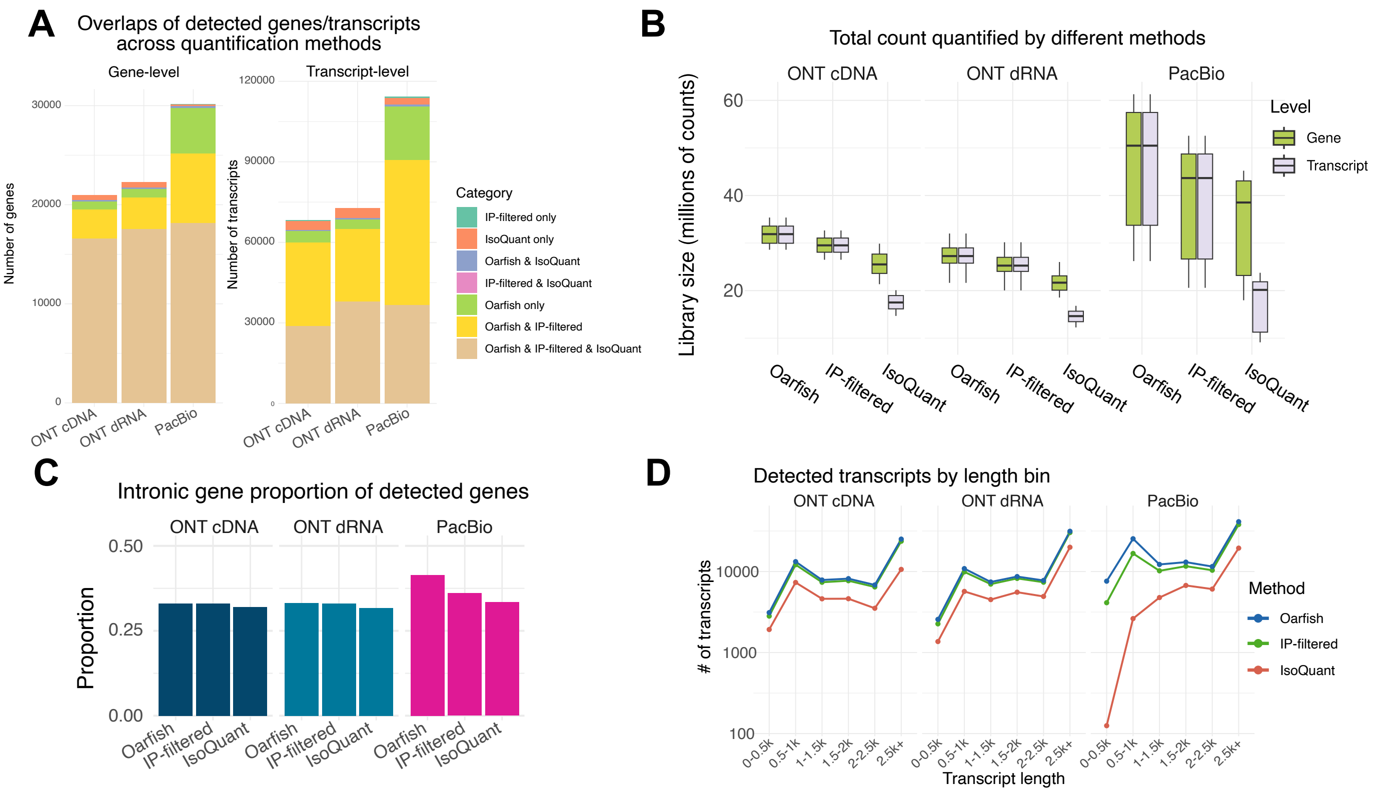

**Supplementary Fig.** 6: **Gene and transcript detection with alternative methods** The gene and transcript detection result depends on the computational workflows. We included alternative workflows in addition to the default one used in the main text. “Oarfish”: the default quantification method used in the main text. “IP-filtered”: an additional filtering step that removes potential reads that are potential produce of internal priming and then uses Oarfish for the quantification and performs gene and transcript detection analysis. “IsoQuant”: Using IsoQuant [(10)] as the quantification tool. **A:** The overlaps of the genes and transcripts detected from 3 different methods. **B:** The total read counts generated from each method. **C:** The proportion of detected genes being “intronic genes”. **D:** The transcript length distribution of the detected transcripts from 3 different methods.

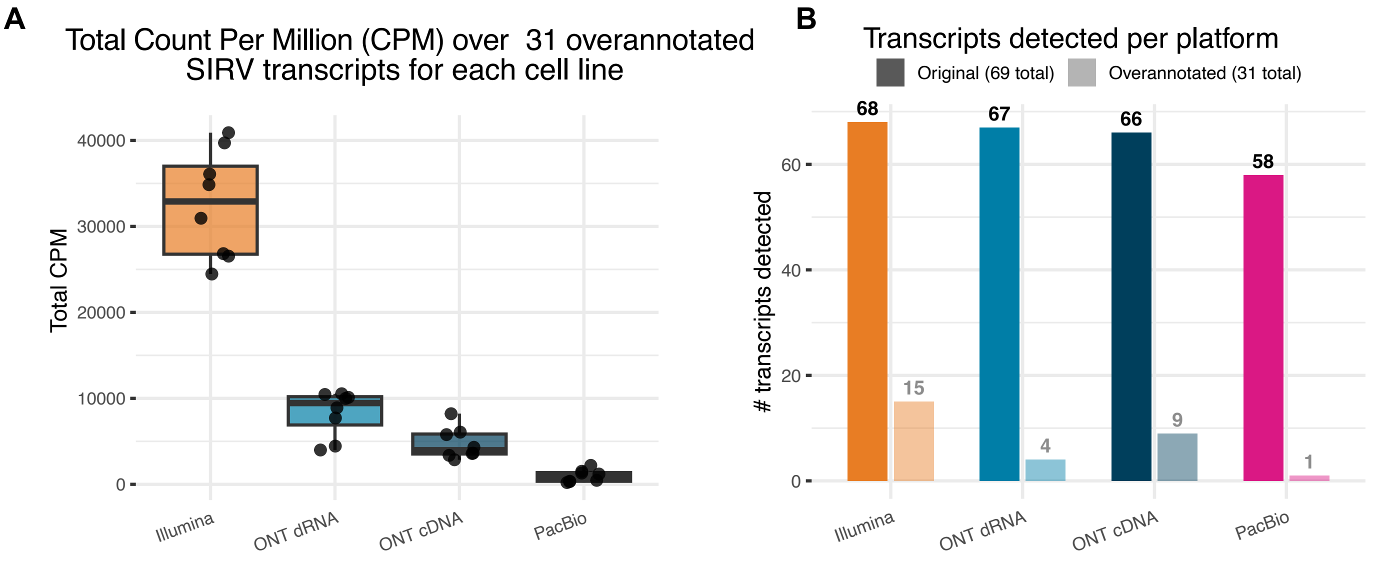

**Supplementary Fig.** 7: **SIRV analysis with over-annotated transcript references:** The official SIRV reference files include an “over-annotated” transcript reference provided by Lexogen. This decoy annotation contains 69 genuine SIRV transcripts (excluding Long SIRVs), together with 31 additional non-existent transcript isoforms. Here, we evaluated transcript detection performance using this over-annotated reference to compare the propensity for false-positive transcript detection across different protocols. **A:** Total counts per million (CPM) assigned to the 31 false-positive SIRV transcripts. Normalisation was performed across all 100 SIRV transcripts. **B:** Number of true and false transcripts identified by each protocol.

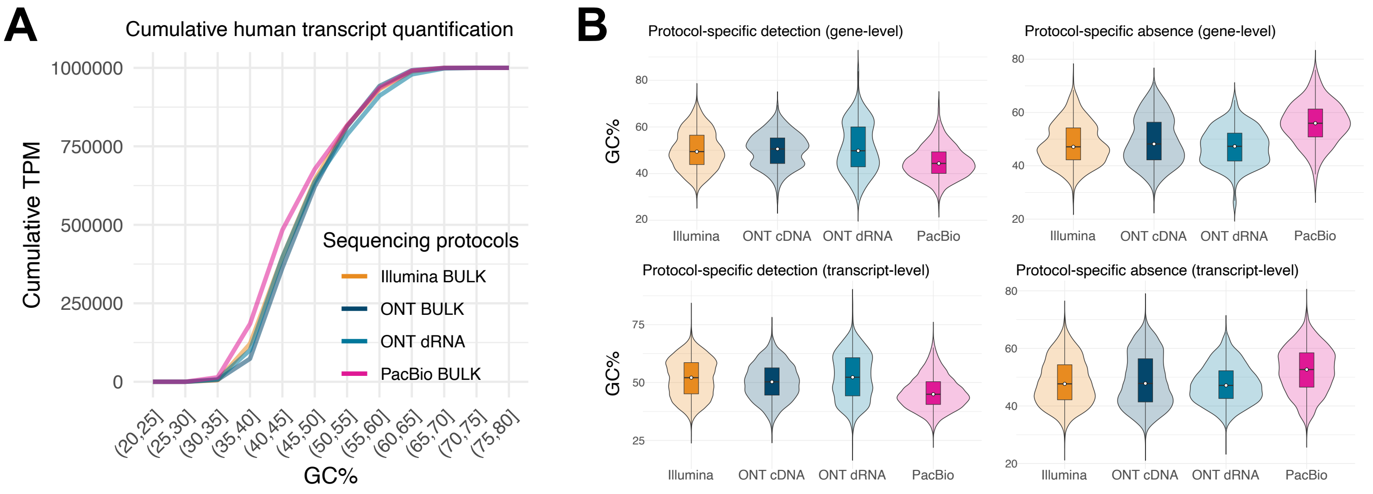

**Supplementary Fig.** 8: **Comparison of GC content across bulk sequencing protocols:**
**A:**: Cumulative transcript abundance (TPM) along transcripts stratified by increasing GC%. **B:** GC% distribution of protocol-specific detection (features detected in one protocol but not in others) and protocol-specific absence (features absent in one protocol but detected by all others). All datasets were downsampled to an identical input size of 20 million reads before transcriptome mapping.

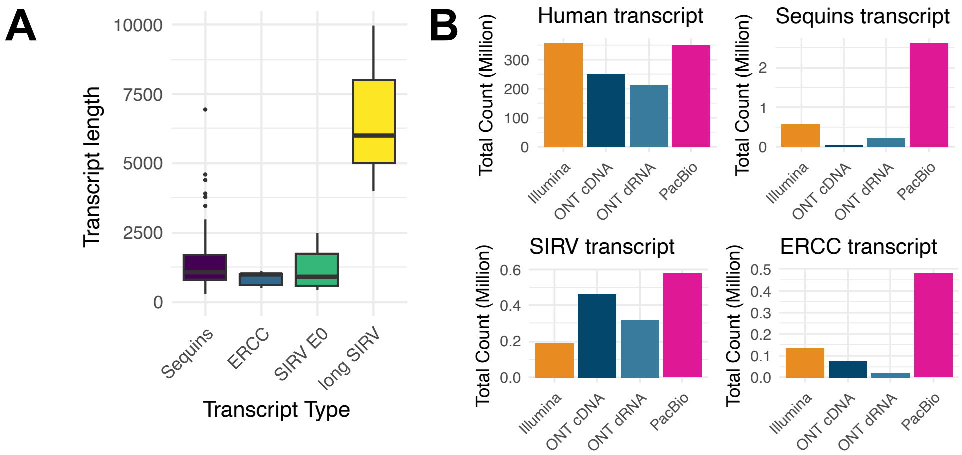

**Supplementary Fig.** 9: **Spike-in transcript overview: A.** Transcript length distribution of the different types of spike-in transcripts. **B.** The total counts assigned to spike-in and human transcripts.

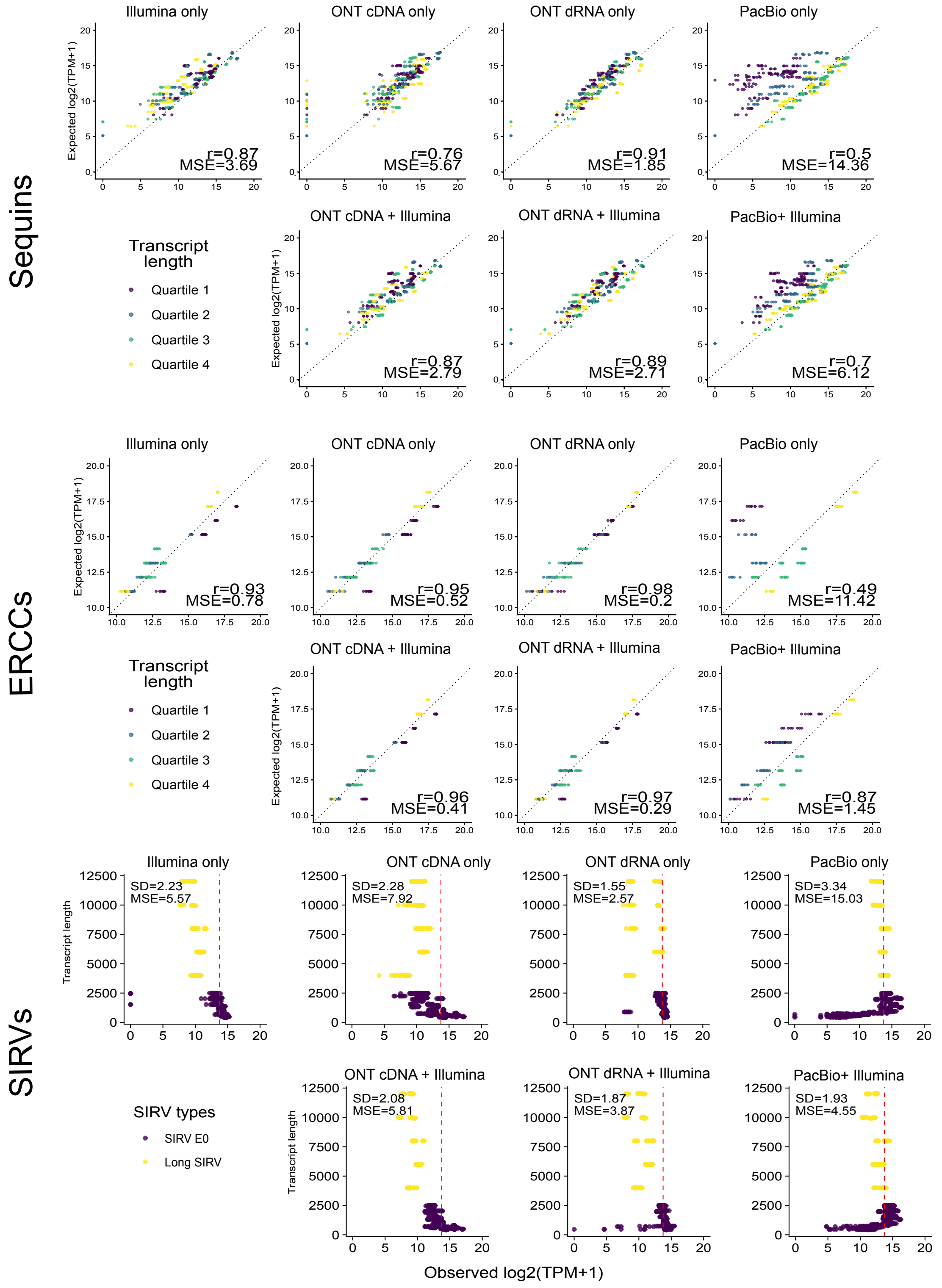

**Supplementary Fig.** 10: **Quantification analysis on spike-in transcripts with alternative long-read quantification tool and hybrid quantification tool:** Comparison of observed $\log_2$(TPM) of spike-in transcripts to expected abundance. Each dot represents one transcript in one cell line, with all eight cell lines shown in each panel. For Sequins and ERCC, each transcript has a distinct expected abundance (y-axis) and is coloured by transcript length. For SIRVs, all transcripts are expected to have equal abundance (indicated by red lines), with transcript length plotted on the y-axis. The long-read-only quantification was performed using IsoQuant (see Methods), and the hybrid quantification was performed using miniQuant [11] (v1.4.1). r: Pearson correlation, MES: Mean square error, SD: Standard deviation

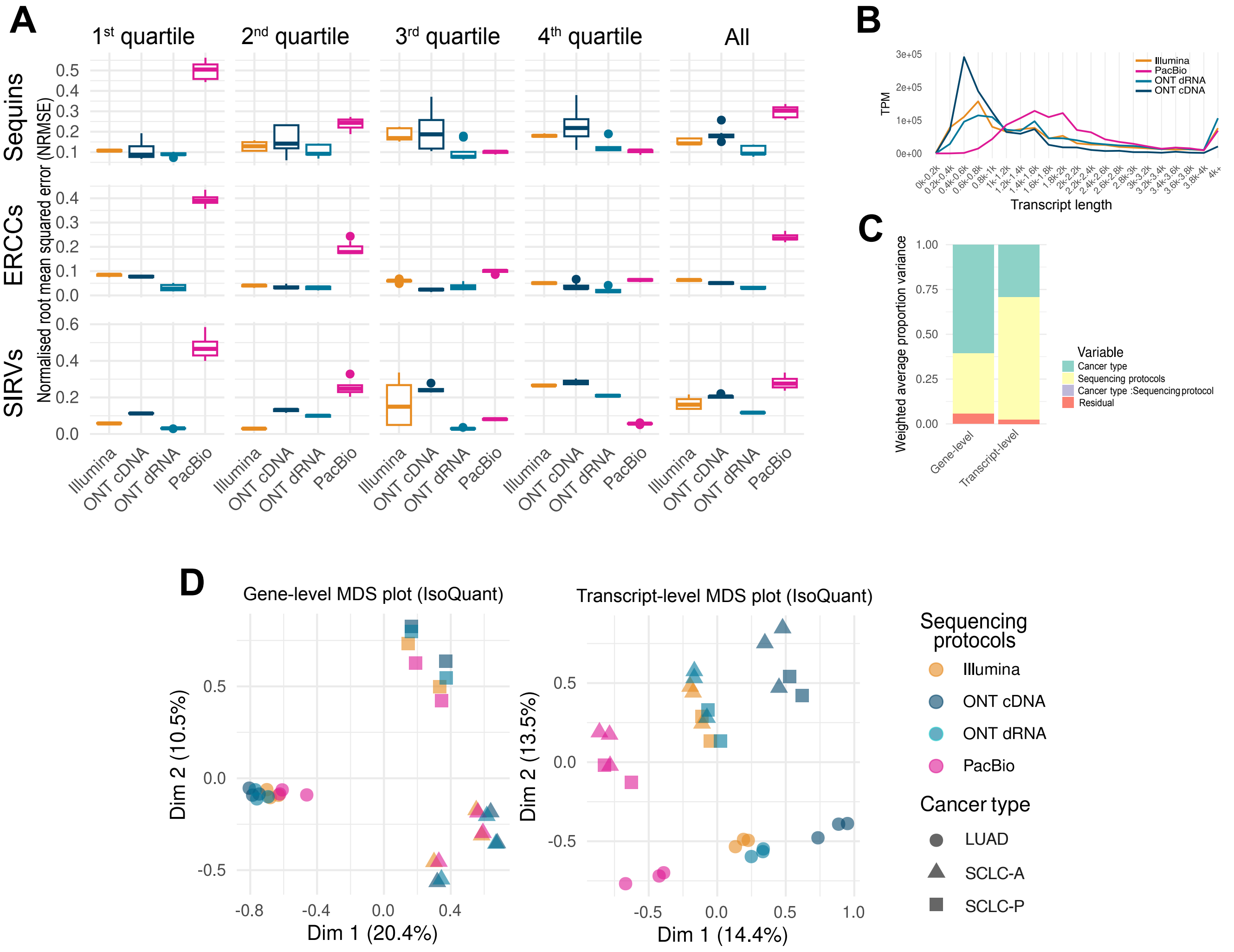

**Supplementary Fig.** 11: **Quantification analysis using IsoQuant:** Unless otherwise specified, all analyses in this manuscript use Oarfish as the default long-read quantification tool. IsoQuant was used instead of Oarfish for this figure to verify that the results are robust to the choice of quantification method. **A.** Normalised root mean squared errors for spike-in transcript quantification in different length quartiles, calculated as: $\text{NRMSE} = \text{RMSE}/\bar{y}$, where $RMSE$ is the root mean square error and $\bar{y}$ represents the mean expected $log_2(TPM + 1)$ of the spike-in transcripts within each respective quartile. **B.** Count distribution across human transcripts with different lengths. **C.** Principal Variance Component Analysis (PVCA) showing the proportion of total variance in gene- and transcript-level quantification explained by each factor and their interactions. Transcript-level counts were summed across eight cell lines and then converted to TPM. **D.** Multidimensional scaling (MDS) plot of samples based on gene-level (D) and transcript-level (E) quantification.

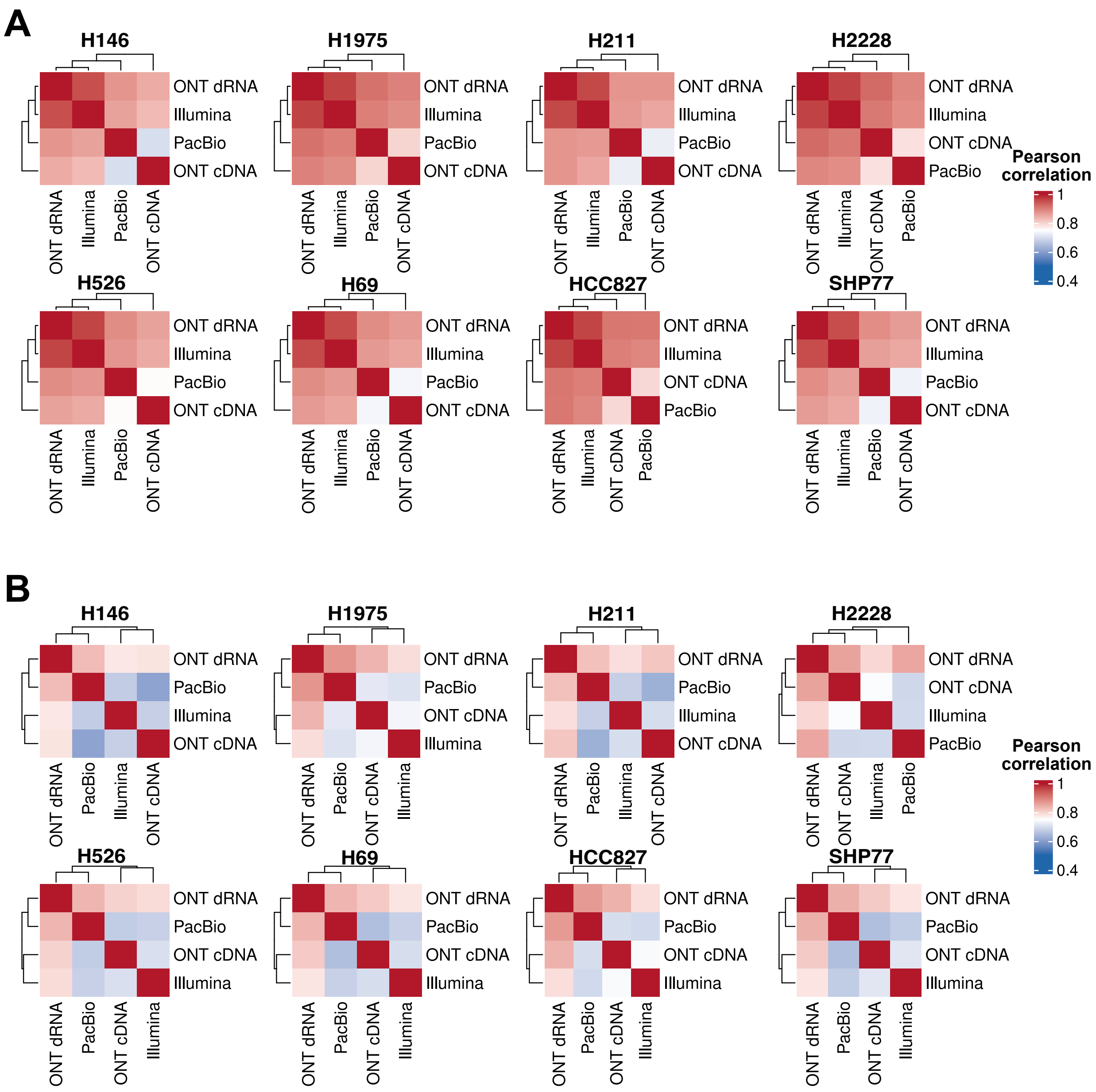

**Supplementary Fig.** 12: **Correlation heatmaps of gene and transcript-level quantification across bulk RNA-seq datasets: A.** Pearson correlation heatmap of gene-level logTPM across bulk RNA-seq datasets. **B.** Pearson correlation heatmap of transcript-level logTPM across bulk RNA-seq datasets.

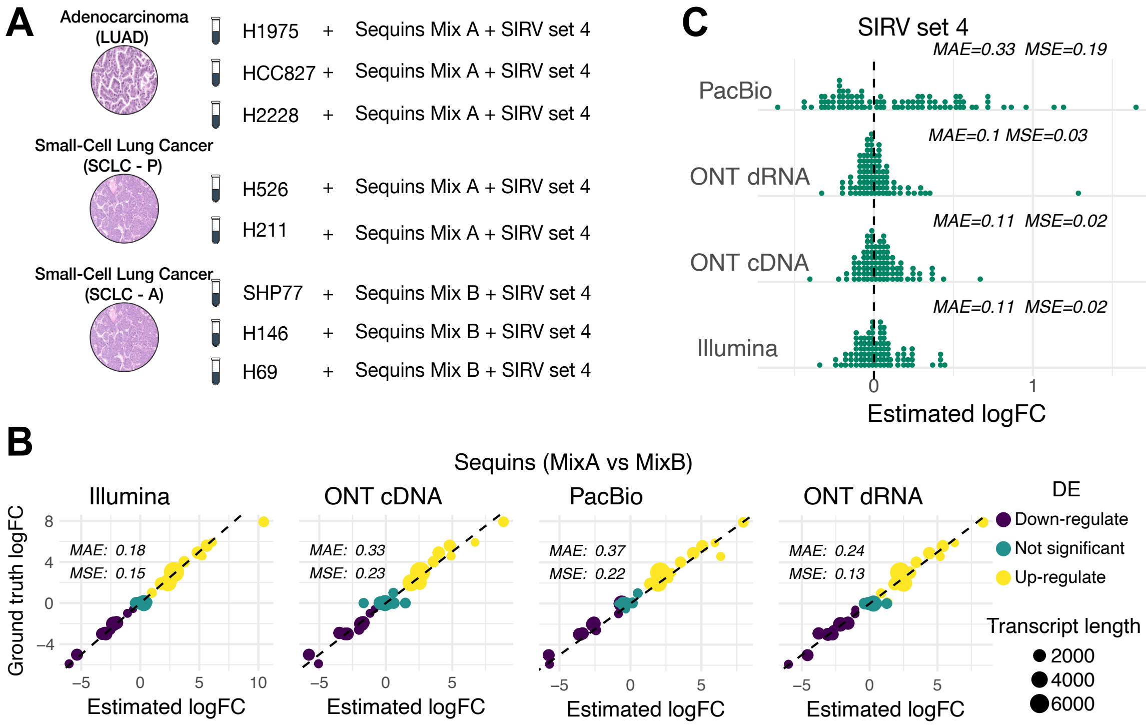

**Supplementary Fig.** 13: **Differential analysis on spike-in transcripts: A.** A summary of different spike-ins added into each sample. **B.** Comparison between the observed log2 fold-change (logFC) and the true logFC in Sequins transcript when comparing samples with Sequins Mix A versus samples with Sequins Mix B added. Each dot is colored by whether it tested to be significant with FDR<0.05 using edgeR. *MSE*: Mean squared error, *MAE*: Mean absolute error. **C.** Distribution of logFC values for SIRV Set 4 transcripts comparing between the same sample groups as B, with an expected logFC of 0.

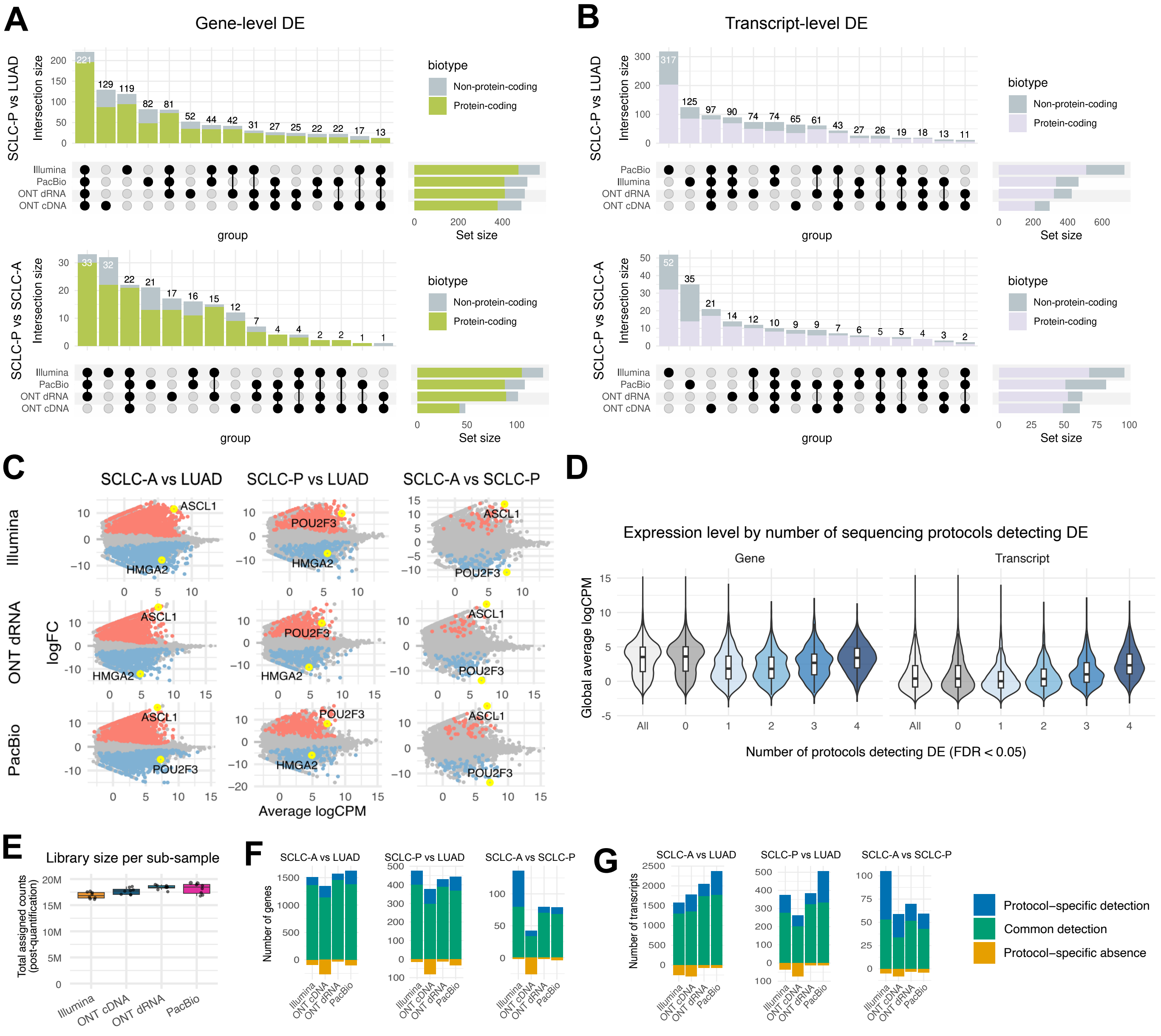

**Supplementary Fig.** 14: **Differential expression analysis of human genes and transcripts on bulk datasets (extended): A,B.** Upset plots showing overlap of differentially expressed (DE) genes (A) and transcripts (B) between comparisons of SCLC-P vs LUAD and SCLC-A vs SCLC-P. **C.** MA plots of gene-level DE analysis for Illumina, PacBio, and ONT dRNA datasets. **D.** Average expression level (logCPM, averaged across all cell lines and protocols) of genes and transcripts stratified by the number of protocols detecting the feature as DE, with "All" representing all tested features. **E–G.** DE analysis from subsampled data (20 million reads per sample), processed using the same pipeline as the full dataset. (D): Estimated library sizes from Oarfish quantification. All datasets were downsampled to an identical input size of 20 million reads before transcriptome mapping. Only minor variations in estimated library size were observed after mapping and quantification. F–G: Number of differentially expressed genes (F) and transcripts (G). DE results are categorised as either common (detected by at least one other platform) or platform-specific. Platform-specific absent signals are shown as negative values, representing DE features identified by all other platforms but missed by the current one.

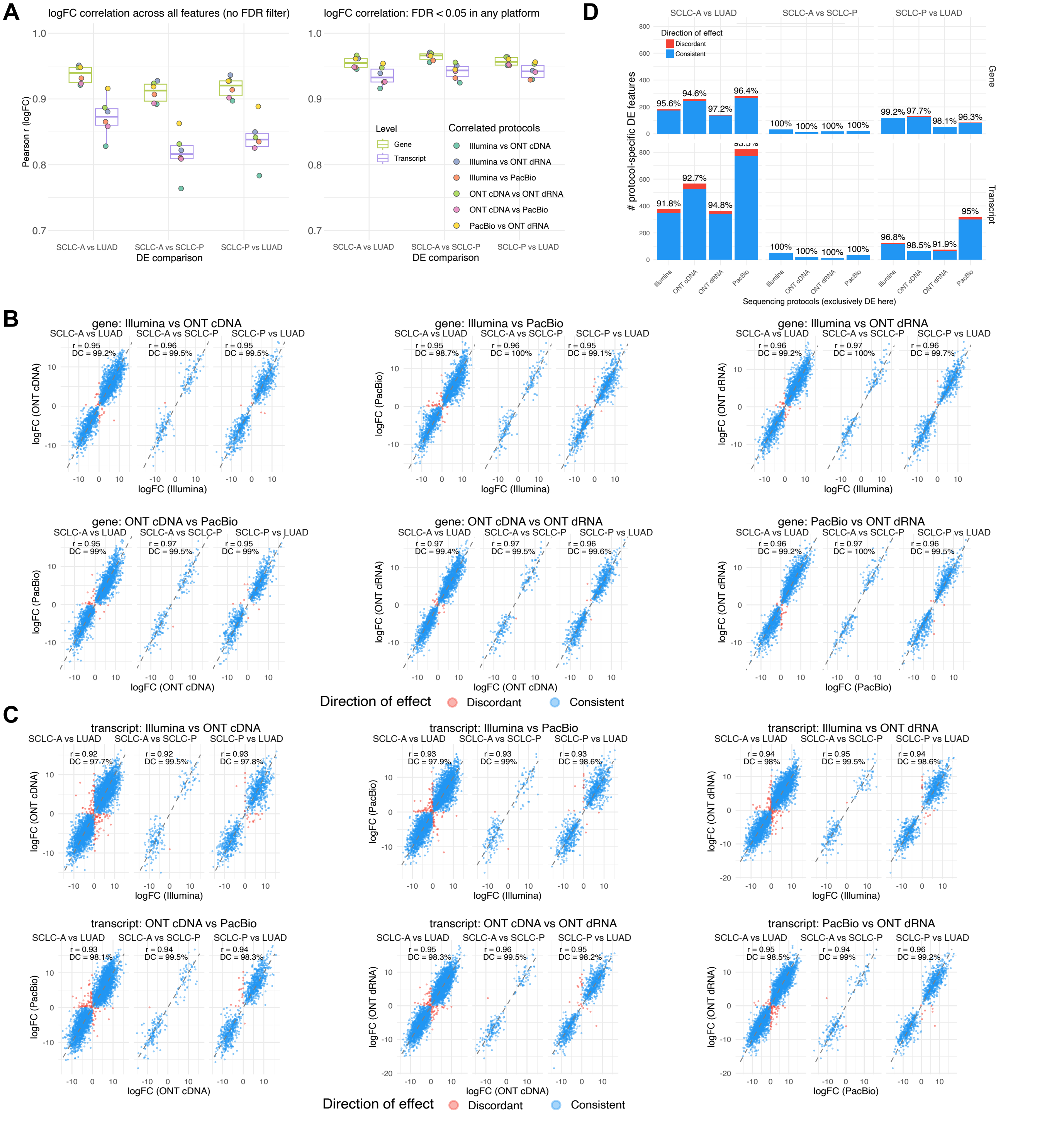

**Supplementary Fig.** 15: **Comparison of log fold change (logFC) of human genes and transcript:** **A.** logFC correlation between pairs of sequencing protocols with all genes and transcripts tested (left) or genes and transcripts reported significant DE in at least one of the protocols. **B,C.** Each panel compares logFC estimates between a pair of sequencing platforms for genes (B) or transcripts (C) identified as significantly differentially expressed (FDR $< 0.05$) in at least one protocol. The dashed line indicates perfect agreement (slope $= 1$). Pearson's correlation coefficient (r) and the proportion of features with concordant effect directions (DC) are shown in each panel, highlighting the consistency of differential expression signals across platforms. **D.** The number and percentage (annotated on the plot) of protocol-specific DE features that were supported by all other protocols with a concordant direction of effect, despite not reaching statistical significance in the other protocols.

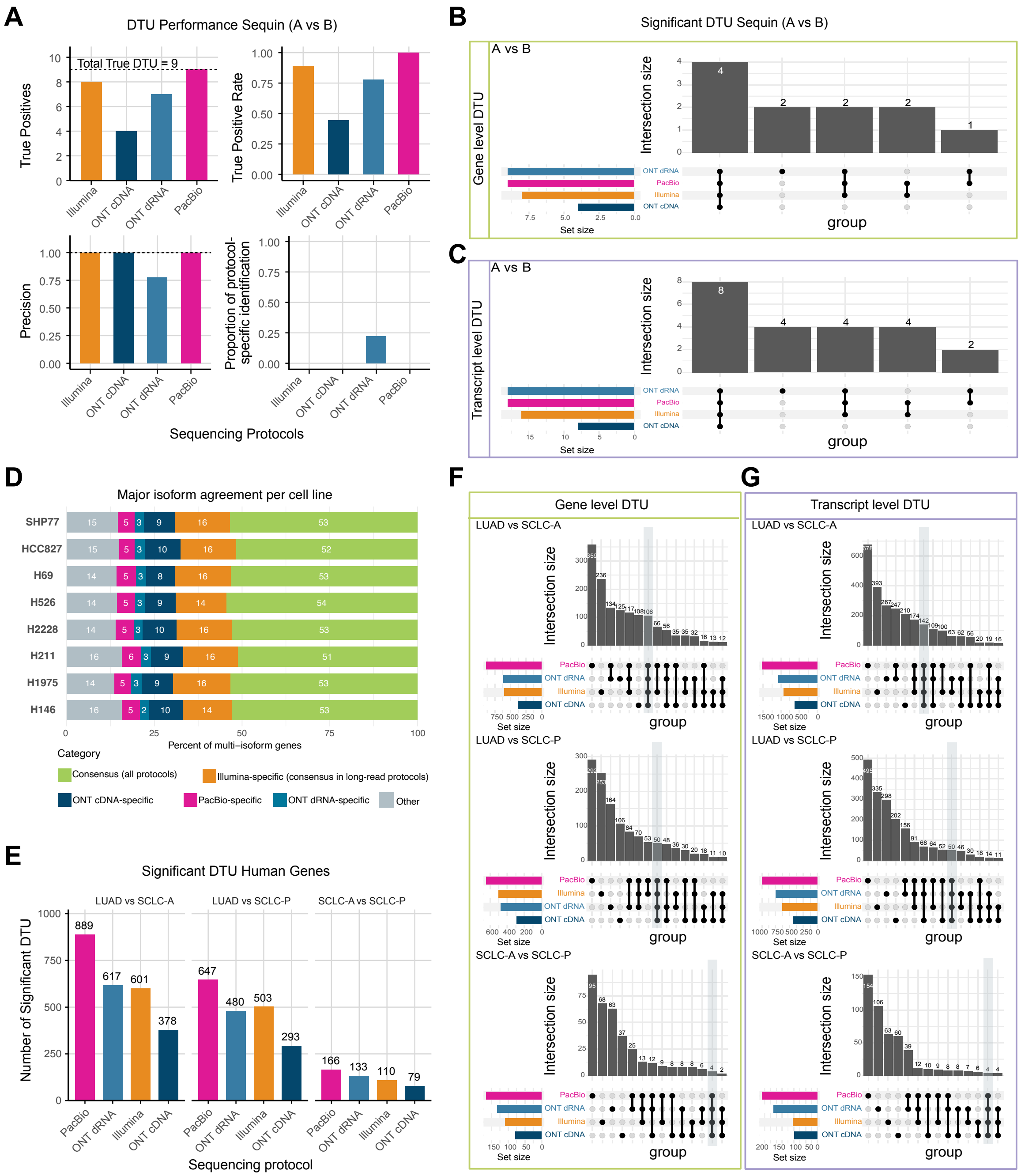

**Supplementary Fig.** 16: **Summary of transcript usage analysis: A–C.** Differential transcript usage (DTU) testing using Sequins transcripts. After filtering for common transcripts and retaining only genes with multiple transcripts, nine genes were known to exhibit DTU between Sequins Mix A and B. A: Summary of gene-level DTU detections. B–C: Overlaps of gene- and transcript-level DTU results across four datasets. **D.** Consistency of major isoform identification in human genes across the four sequencing protocols. For each protocol, the major isoform was defined as the transcript with the highest count, after applying the same filtering criteria used in the DTU analysis to restrict to genes with multiple transcripts. **E.** Total number of significant DTU genes identified per sequencing protocol, based on pairwise comparisons across the three cancer types. **F–G.** Overlaps of gene- and transcript-level DTU results from human genes across the four sequencing protocols.

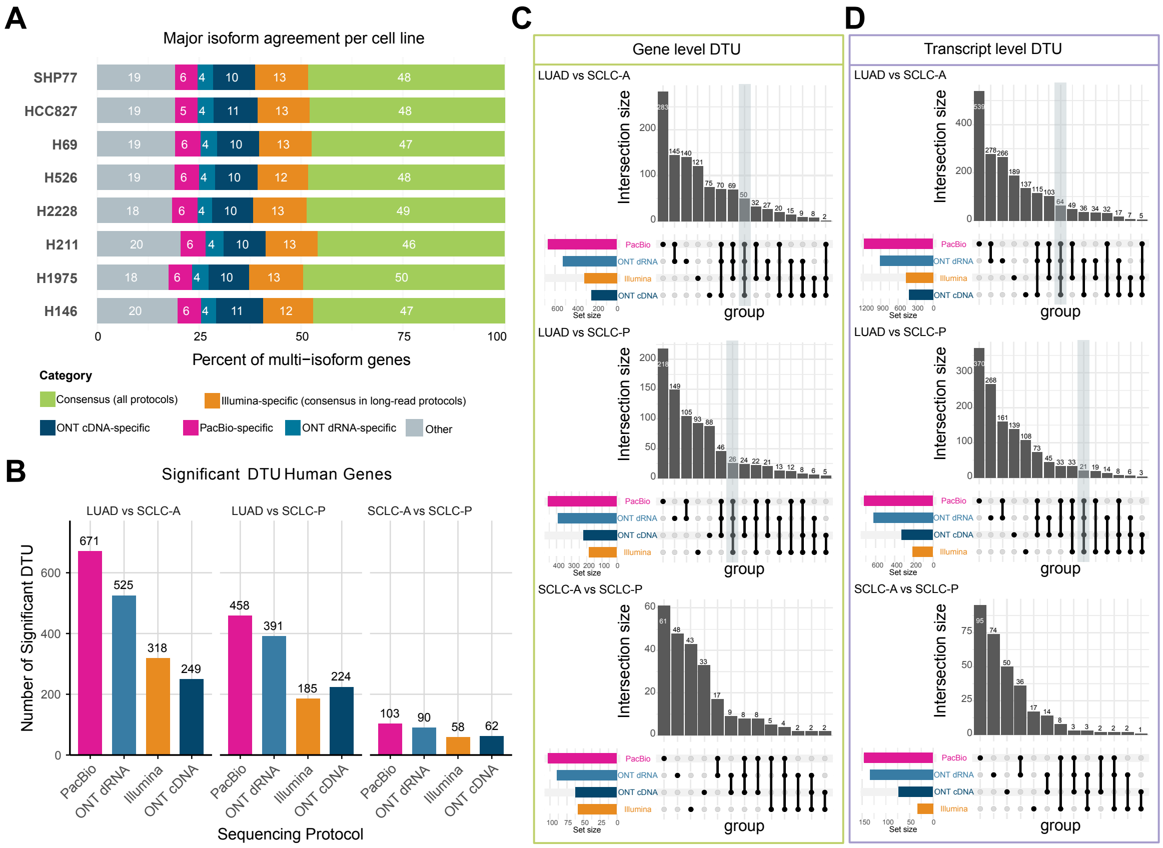

**Supplementary Fig.** 17: **Summary of transcript usage analysis on down-sampled data.** All cell line samples were down-sampled to 20M reads prior to quantification and DTU analysis. **A.** Consistency of major isoform identification in human genes across the four sequencing protocols. For each protocol, the major isoform was defined as the transcript with the highest count, after applying the same filtering criteria used in the DTU analysis to restrict to genes with multiple transcripts. **B.** Total number of significant DTU genes identified per sequencing protocol, based on pairwise comparisons across the three cancer types. **C–D.** Overlaps of gene- and transcript-level DTU results from human genes across the four sequencing protocols.

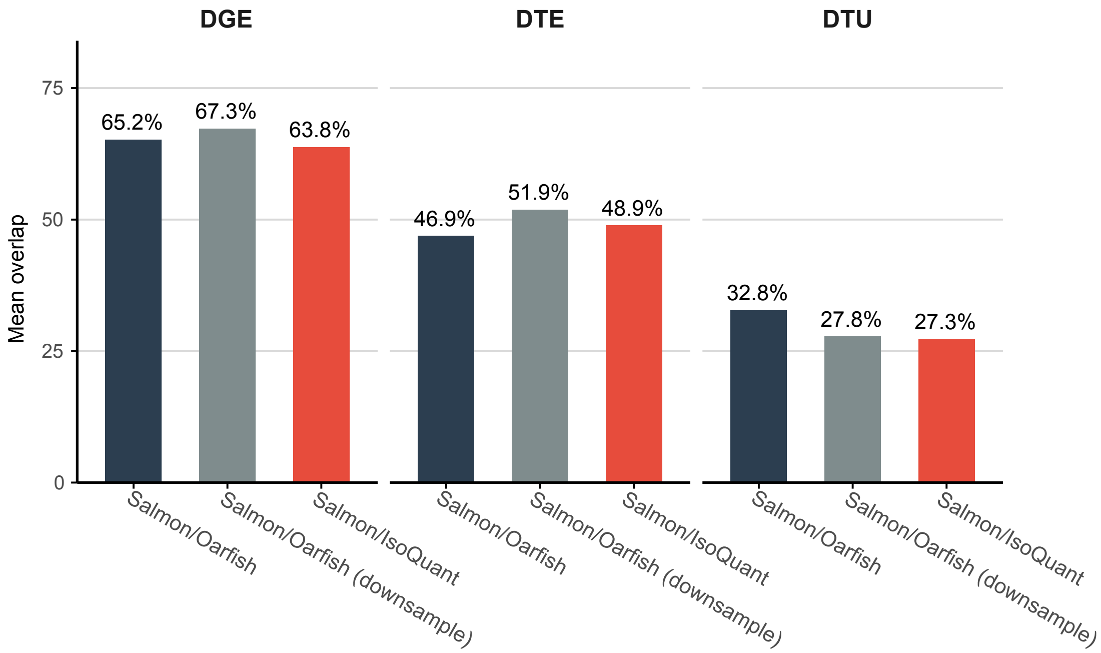

**Supplementary Fig.** 18: **Summary of across-protocol concordance of differential analysis:** The mean overlap (see Methods) of differential gene expression (DGE), differential transcript expression (DTE), and differential transcript usage (DTU) across the four sequencing protocols. Concordance progressively decreased from DGE to DTE and DTU. Results from two long-read quantification tools (Oarfish and IsoQuant), as well as analyses using downsampled input data (20M reads per cell line for each sequencing protocol), were included to assess the robustness of this trend across quantification methods and sequencing depths. For DTU, the concordance shown here represents gene-level overlap, i.e., overlap of genes identified as exhibiting DTU.

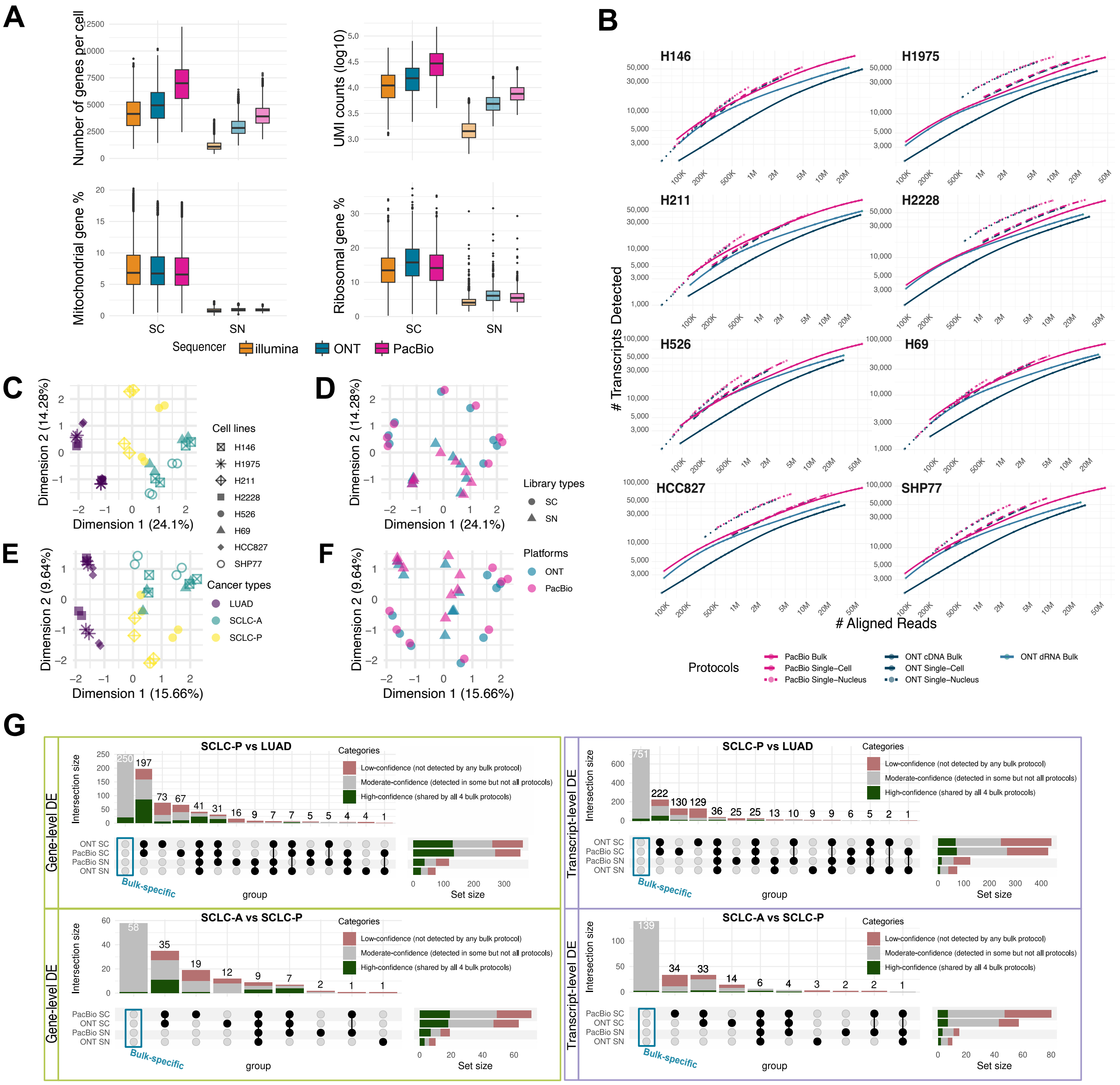

**Supplementary Fig.** 19: **Analysis of single-cell and single-nucleus datasets (extended): A.** Single-cell quality control plots. No major quality issues were observed. As expected, single-nucleus libraries showed lower mitochondrial and ribosomal gene content due to the removal of cytoplasmic components. **B.** Comparing the number of transcripts detected in each bulk or pseudobulk sample (defined as transcripts with a minimum count of 5). **C-F.** Gene-level and transcript MDS plots of pseudobulk samples. **G.** Gene- and transcript-level pseudobulk DE results comparing SCLC-P vs LUAD and SCLC-A vs SCLC-P.

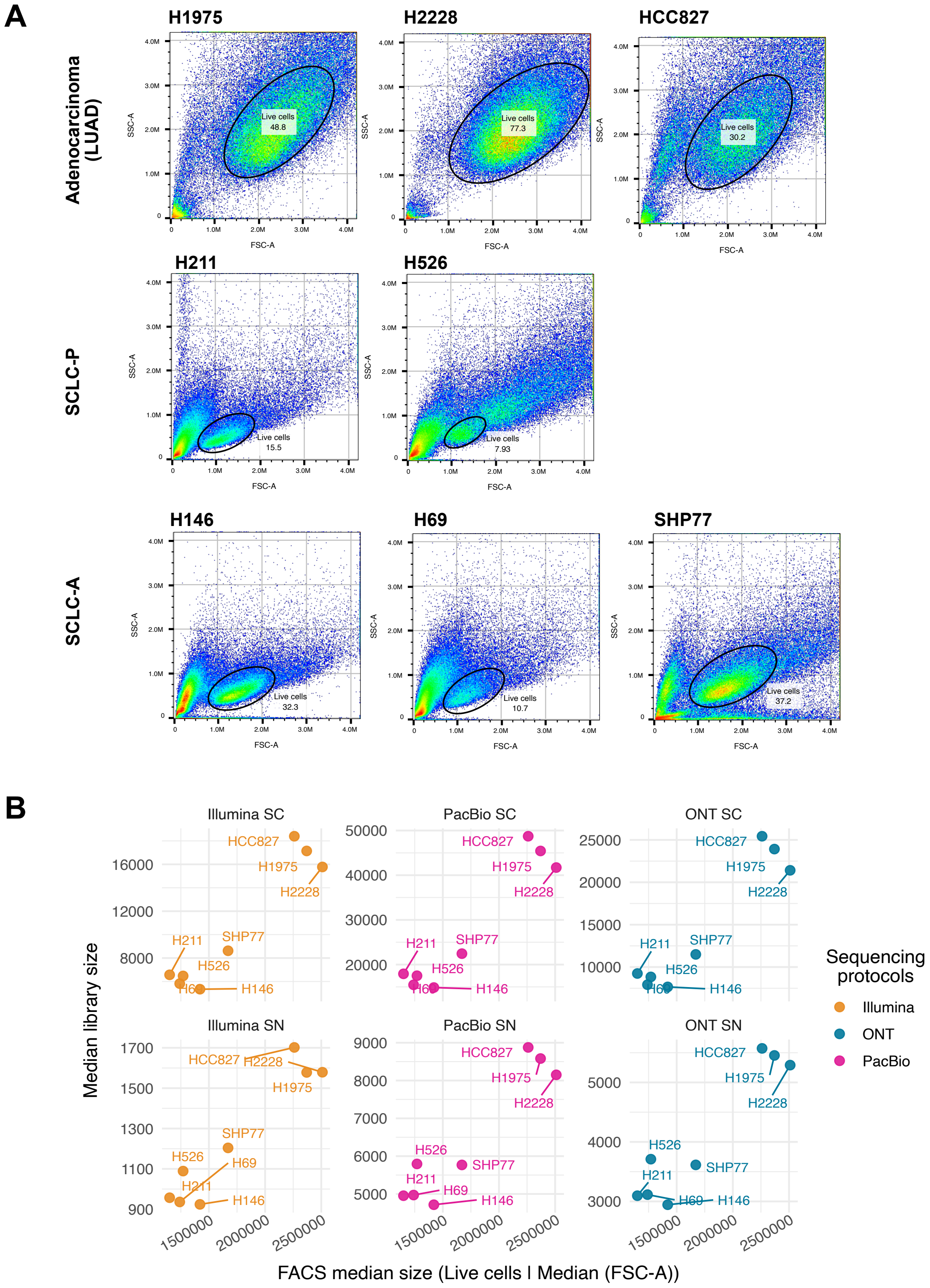

**Supplementary Fig.** 20: **Cell size analysis using flow cytometry: A:** Flow cytometry plots showing live cell distribution using constant forward scatter (20V) and side scatter (55V) voltages. Live cells gated based on DAPI exclusion. **B:** Scatter plot of Forward Scatter Area (FSC-A) of live cells versus the single-cell/single-nucleus library size in the long-read datasets.

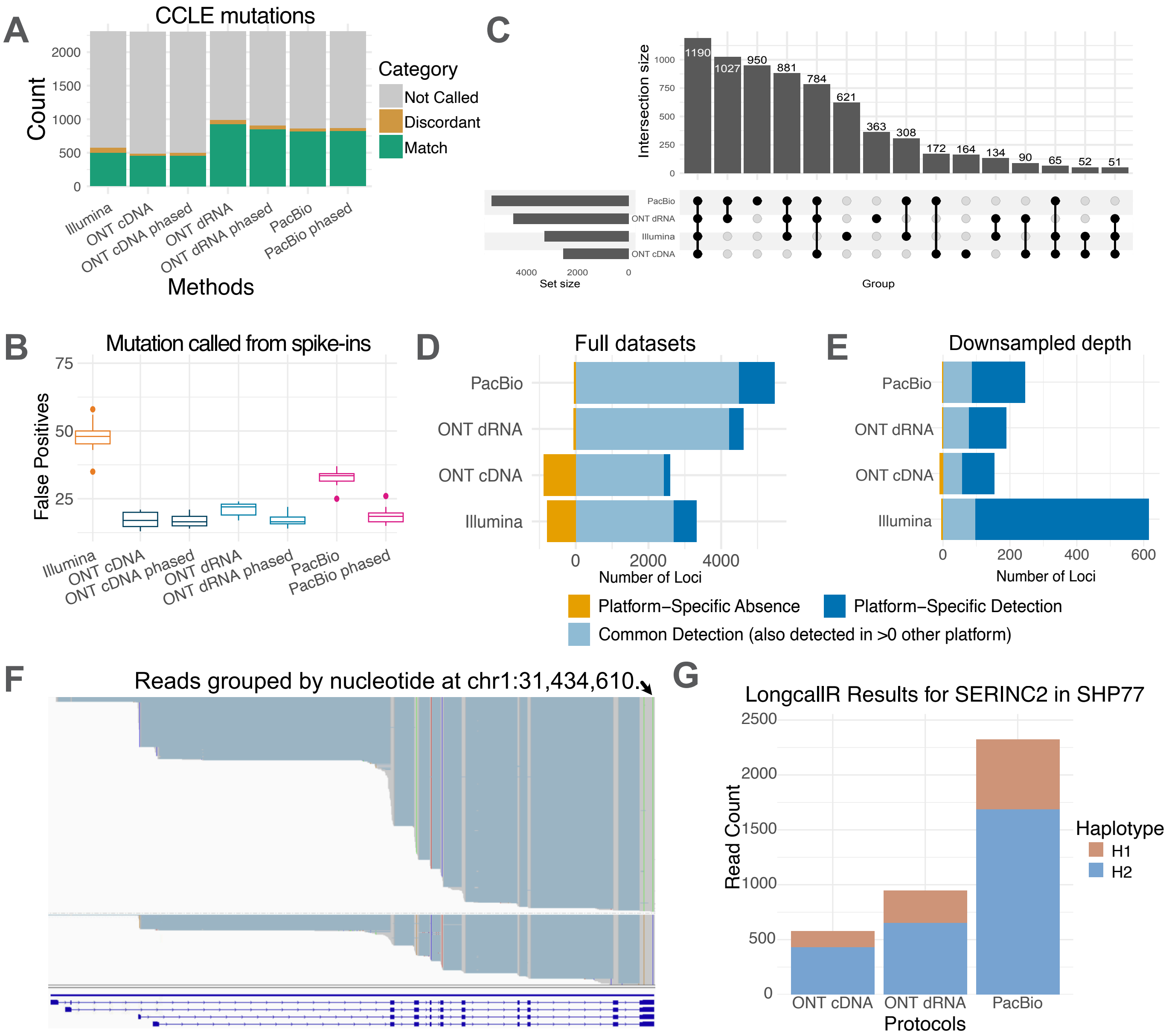

**Supplementary Fig.** 21: **Variance calling and allelic imbalance analysis (extended): A.** Overlap of variants identified by multiple methods with known CCLE somatic mutations. In Clair3-RNA, we compared variant calling with the phasing model enabled versus disabled to evaluate whether phasing improves calling accuracy. **B.** The variants called on spike-ins using the same methods as in A **C.** The overlaps of allelic imbalance events identified from different bulk RNAseq datasets. The allelic imbalance was assessed using a two-sided binomial test ($H_0$: alternative allele frequency $= 0.5$), with significance determined at BH-adjusted p-value $< 0.05$. **D.** A summary of platform-specific allelic imbalance identification **E.** To enable fair comparison of allelic imbalance detection across platforms with varying sequencing depths, we downsampled the read coverage at each locus to 30 reads. This standardisation controls for differences in statistical power and ensures platform-specific differences reflect true biological or technical variation rather than depth-related bias. **F-G.** We illustrate allele-specific expression (ASE) of the SERINC2 gene in the SHP77 cell line. A representative Integrative Genomics Viewer (IGV) screenshot from the bulk PacBio dataset (F) highlights distinct haplotypes, with reads grouped by the nucleotide at chr1:31,434,610 and showing clear phasing with other exonic variants. The corresponding barplot (G) quantifies the allelic expression imbalance from all long-read protocols using LongcallR[(12)].

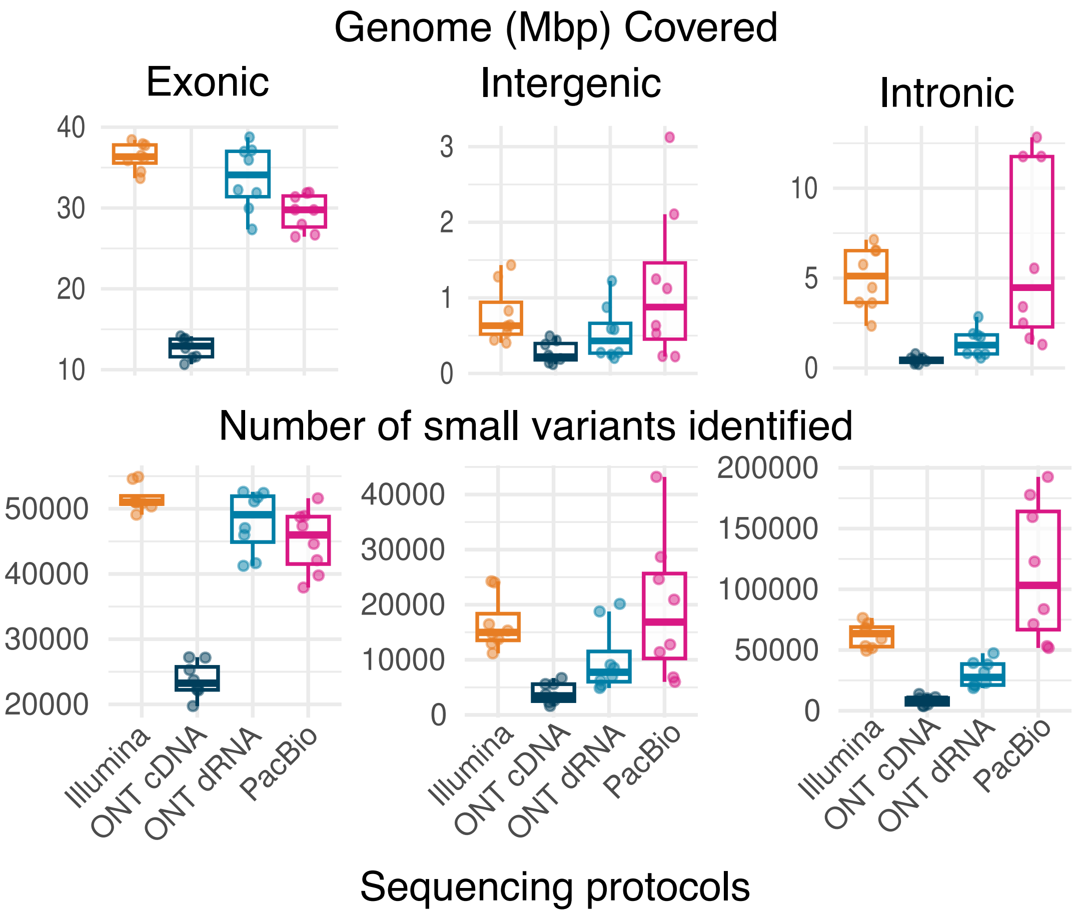

**Supplementary Fig.** 22: **Small variant analysis with gigabase-downsampled data (extended):** Coverage and number of variants detected across different genomic regions. To ensure a fair comparison across sequencing protocols, the input long-read datasets were downsampled to match the total gigabases of the short-read data (see Methods). A minimum read depth of 30 was required for a genomic position to be considered covered.

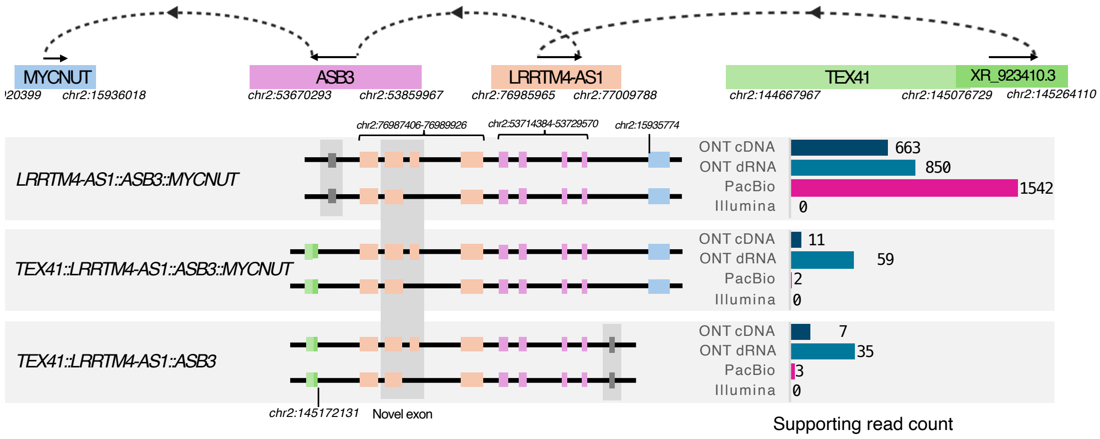

**Supplementary Fig.** 23: **Example of detected four-gene fusion:** Illustration of a four-gene fusion event involving TEX41, MYCNUT, ASB3, and LRRTM4-AS1, detected concordantly across all three long-read protocols in the H69 cell line.

### Supplementary Information

#### 1.1 Summary of Spike-ins

The Sequins Mix A and B contain identical transcript isoforms at different relative abundances, providing a ground truth for differential expression analyses. The SIRV-Set 4 comprises three components: the SIRV isoform mix (E0), the ERCC mix, and the long SIRVs. The E0 mix contains 69 isoforms at equal molar concentrations, serving as a control for isoform complexity. The long SIRVs include 16 non-isoform transcripts ranging from 4 to 12 kb, also at equal molar concentrations as the E0 mix. The ERCC mix consists of 92 non-isoform transcripts spanning a dynamic concentration range, enabling quantification assessment across a wide expression spectrum. Together, these spike-ins provide a robust ground truth for systematically benchmarking sequencing protocols (Supplementary Table 1).
